# Uncovering molecular grammars of intrinsically disordered regions that organize nucleolar fibrillar centers

**DOI:** 10.1101/2022.11.05.515292

**Authors:** Matthew R. King, Andrew Z. Lin, Kiersten M. Ruff, Mina Farag, Wei Ouyang, Michael D. Vahey, Emma Lundberg, Rohit V. Pappu

## Abstract

The nucleolus is a multilayered structure. Each layer is thought to be a compositionally distinct phase, although how these phases form and interface with one another remains unclear. Using computational, proteomics, *in vitro*, and *in vivo* studies, we uncover distinct molecular grammars within intrinsically disordered regions (IDRs) of nucleolar proteins that localize to fibrillar centers (FCs) and dense fibrillar components (DFCs). FC and DFC proteins feature two distinct types of IDRs namely those with long tracts of acidic residues and those with blocks of lysines interspersed by acid-rich-regions. We find that phase separation driven by complex coacervation in mixtures of nucleolar proteins, with their distinctive IDRs, and ribosomal DNA and RNA molecules is sufficient to drive the formation of structural facsimiles of FCs and DFCs.

**One-Sentence Summary:** Facsimiles of core nucleolar substructures were reconstituted via phase separation of key protein and nucleic acid mixtures.

## Main Text

The cellular nucleus is the site of multiple coexisting membraneless biomolecular condensates that consolidate discrete factors for designated functions ^1^. Biomolecular condensates are thought to form via coupled associative and segregative phase transitions ^2^, which we refer to hereafter as condensation ^3^. The largest condensate is the nucleolus featuring a hierarchical organization is directly relevant for enabling the assembly-line process of ribosomal biogenesis ^4,5^. Ribosomal DNA (rDNA) and the transcriptional machinery are housed in the inner-most layer, the fibrillar center (FC). The transcribed premature ribosomal RNA (pre-rRNA), is folded, matured, and complexed with ribosomal proteins in the dense fibrillar component (DFC) and the granular component (GC) (Fig. SI1). These layers, which have long been appreciated by electron microscopy ^6^, are thought to be coexisting phases ^4,5,7^. Despite this, our knowledge of how constituent proteins and nucleic acids drive condensation and hierarchical organization of the coexisting phases is far from complete.

Proteins that drive condensation generally harbor multiple substrate-binding domains and specific types of intrinsically disordered regions (IDRs) ^8^. Drivers of distinct condensates have distinct molecular grammars^9–11^. The framework of stickers-and-spacers^8^ has proven to be useful for describing molecular grammars because it partitions protein sequences into cohesive stickers and solubility-determining spacers ^8^. Associative interactions that provide part of the drive for condensation are governed by the numbers of stickers, the different sticker types, and the molecular topologies of stickers vs. spacers. These features enable many distinct condensates to assemble and coexist within a cell. Evidence for this comes from recent studies on stress granules, nuclear speckles, and unfolded protein deposits ^9,10,12^. Establishing the molecular grammars that drive and regulate condensation transitions of different bodies is therefore key to understanding how substrate binding domains and IDRs combine to encode sequence-specific driving forces for condensate formation and for the regulation of their compositions.

To identify condensate-specific scaffolds and condensate-driving molecular grammars within IDRs, we developed a computational pipeline that combines comparisons of compositional features and alignment-free z-scores of various binary sequence patterns ^13,14^. We profiled 1,156 IDRs across five distinct cellular condensates that have been inventoried by cellular proteomics (Figs 1A, SI2, and methods) ^15–17^. Our analysis returned sequences with known molecular grammars. This includes the finding that stress granule IDRs feature high fractions of aromatic residues (Fig. 1A, grayed boxes) ^9,11^. Interestingly, we found that nucleolar IDRs had the largest number of distinctive sequence features with *p*-values < 10^−5^ and the highest single enrichment *p*-value for a sequence feature. Among the distinctive features was the presence of IDRs with blocks of like-charged residues (Fig. 1A). This is consistent with previous observations ^18,19^ and is replicated in unbiased hierarchical clustering of sequence features of all nucleolar IDRs (Fig. SI 3 and SI 4A). Among the distinctive nucleolar IDRs are those featuring high but roughly equal fractions of lysines and glutamic acids and segregation of oppositely charged residues into distinct clusters along the linear sequence (Figs. 1B and SI 4B). Accordingly, proteins with these IDRs are representatives of a molecular grammar comprising K-blocks interspersed by E-rich regions (referred to hereafter as K-blocks+ERRs). Ly-1 antibody reactive (LYAR) is an example of this class (Fig. 1B). This protein is thought to have scaffolding and chaperoning functions in nucleoli ^20^. The second distinctive molecular grammar is the presence of polyelectrolytic IDRs characterized by long tracts rich in Asp and Glu; these we refer to as D/E-tracts. The nucleolar biogenesis protein Upstream Binding Transcription Factor (UBF) belongs to this class (Fig. 1B), which also includes key nucleolar proteins such as nucleophosmin (NPM1) and nucleolin (NCL).

**Fig 1:**
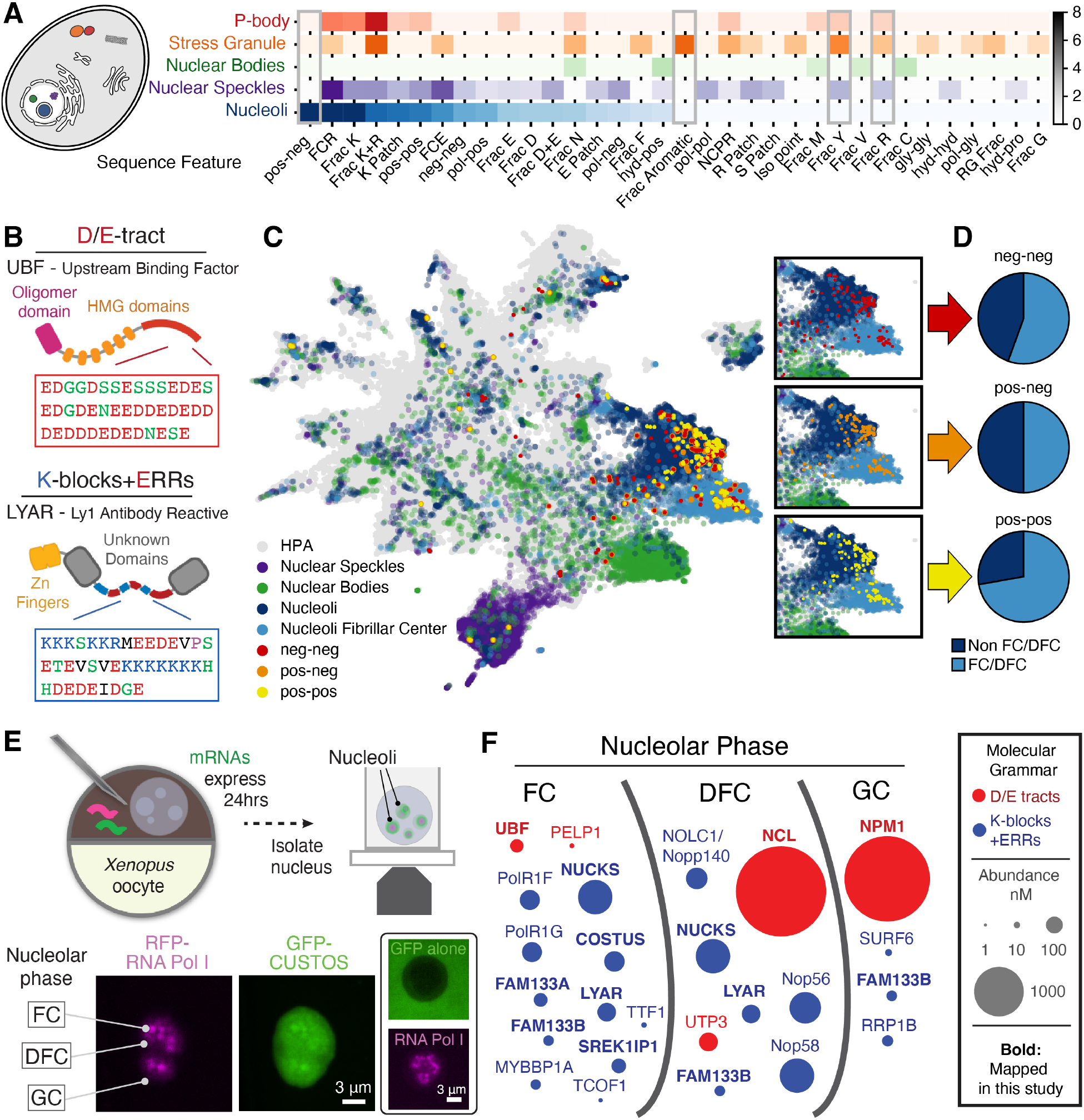
The FC and DFC phases of the nucleolus harbor proteins with K-blocks+ERRs and D/E-tract IDRs. (**A**) Enriched sequence features of IDRs for proteins from five biomolecular condensates. Sequence features come in two categories: the patterning of pairs of amino acid types (x-x) or composition (see methods). Only sequence features with which at least one condensate had a p-value < 0.05 are shown. Scale bar denotes the signed log_10_ (*p*-value) when compared to the remaining human IDRome. Positive signed log_10_ (*p*-value) values imply that feature is more enriched (composition) or blockier (pattern) in each condensate when compared to the rest of the human IDRome (see methods). (**B**) Example nucleolar proteins containing D/E-tract and K-blocks+ERRs within IDRs. (**C**) UMAP representation of the Subcellular Atlas of the Human Protein Atlas with proteins highlighted by subcellular location or high scoring patterning features that involve charged residues (see methods). Here, the locations of 18 nucleolar IDRs with highest z-scores quantifying the patterning of charged residues (neg-neg, pos-neg, and neg-neg) are shown. For nucleolar proteins, a high scoring pos-pos z-score value generally implies the IDR contains K-blocks+ERRs. Likewise, a high scoring pos-neg or neg-neg z-score value generally implies the IDR contains either K-blocks+ERRs or a D/E-tract depending on its net charge per residue. Insets highlight the preference for these highest scoring sequences to be localized to nucleoli. (**D**) Fraction of the sequence with highest-scoring z-scores for neg-neg, pos-neg, and pos-pos patterns that are localized to the FC/DFC phases. (**E**) Schematic of method for obtaining and imaging living nucleoli (see Methods for details). Representative image of a nucleolus expressing GFP-tagged CUSTOS/c12orf43 and RFP-tagged PolRIe subunit of RNA Polymerase I; inset shows a nucleolus expressing GFP alone, as a negative control. GC, DFC, and FC phases are annotated; scale bars = 3 μm. (**F**) Spatial map of proteins with high-scoring K-blocks+ERRs and D/E-tracts in the FC/DFC/GC phases of the nucleolus.

To determine the roles of the proteins with IDRs that have K-blocks+ERRs and /or D/E-tracts, we first identified their localizations within nucleoli. Plotting the locations of these proteins on the uniform manifold approximation and projection for dimension reduction (UMAP) representation of the Subcellular Atlas of the Human Protein Atlas, we found that they were disproportionately enriched in regions that correspond to the FC and DFC (Fig. 1 C/D). To gain precision regarding the localization to the FC vs. DFC, we imaged previously unmapped proteins using living *Xenopus laevis* germinal vesicles obtained from oocytes expressing green or red fluorescent protein (FP)-tagged proteins of interest from microinjected mRNAs. While FPs alone are partially excluded from nucleoli, all proteins with high scoring K-blocks+ERRs and D/E-tract IDRs when fused to FPs readily localized to nucleoli (Figs. 1E and SI5A-D). In contrast, FP-tagged versions of the N-terminal IDR of Huntingtin (Httex1) were excluded to the same extent as free FPs (Fig. SI5E-H). Httex1 consists of an extended polyglutamine tract, and has been shown to drive the formation of distinct cellular condensates ^10^. The exclusion of FP-tagged Httex1 from nucleoli and the strong localization of FP-tagged proteins harboring IDRs with high-scoring K-blocks+ERRs or D/E-tracts highlights the role of molecular grammar in determining specificity of localization. Importantly, we find that proteins harboring IDRs with high-scoring K-blocks+ERRs or D/E-tracts are enriched in the FC and DFC regions (Fig. 1F). However, NCL is preferentially excluded from the FC, whereas UBF is preferentially excluded from the DFC and GC. These localization biases, the scaffold-like domain architectures, and the presence of unique sequence biases for FC/DFC proteins led us to hypothesize that proteins featuring IDRs with either of the two categories of molecular grammars are likely to be key determinants of the condensation and organization of the FC and DFC.

Overexpressing a protein that directly contributes to condensation often enlarges the constituent biomolecular condensate. This has been shown for overexpression of NPM1, which leads to enlarged nucleolar GCs ^7^. We carried out a targeted overexpression screen of FC factors identified in our multipronged analyses. Overexpression of UBF resulted in significant FC enlargement (Figs. 2A-B; Fig. SI6A/C). This is consistent with UBF knockdown leading to FC loss in cell culture among other similar observations that collectively implicate UBF as a key FC scaffold ^21^. Of the other proteins tested, only LYAR overexpression resulted in FC enlargement. Interestingly, overexpression of LYAR, CUSTOS, or SREK1IP1 also led to the formation of *de novo* condensates. This behavior is reminiscent of a phenotype that has been observed upon overexpression of the DFC protein, fibrillarin (Figs. 2A/C, SI5C-D, SI6B-C) ^7^.

**Fig 2:**
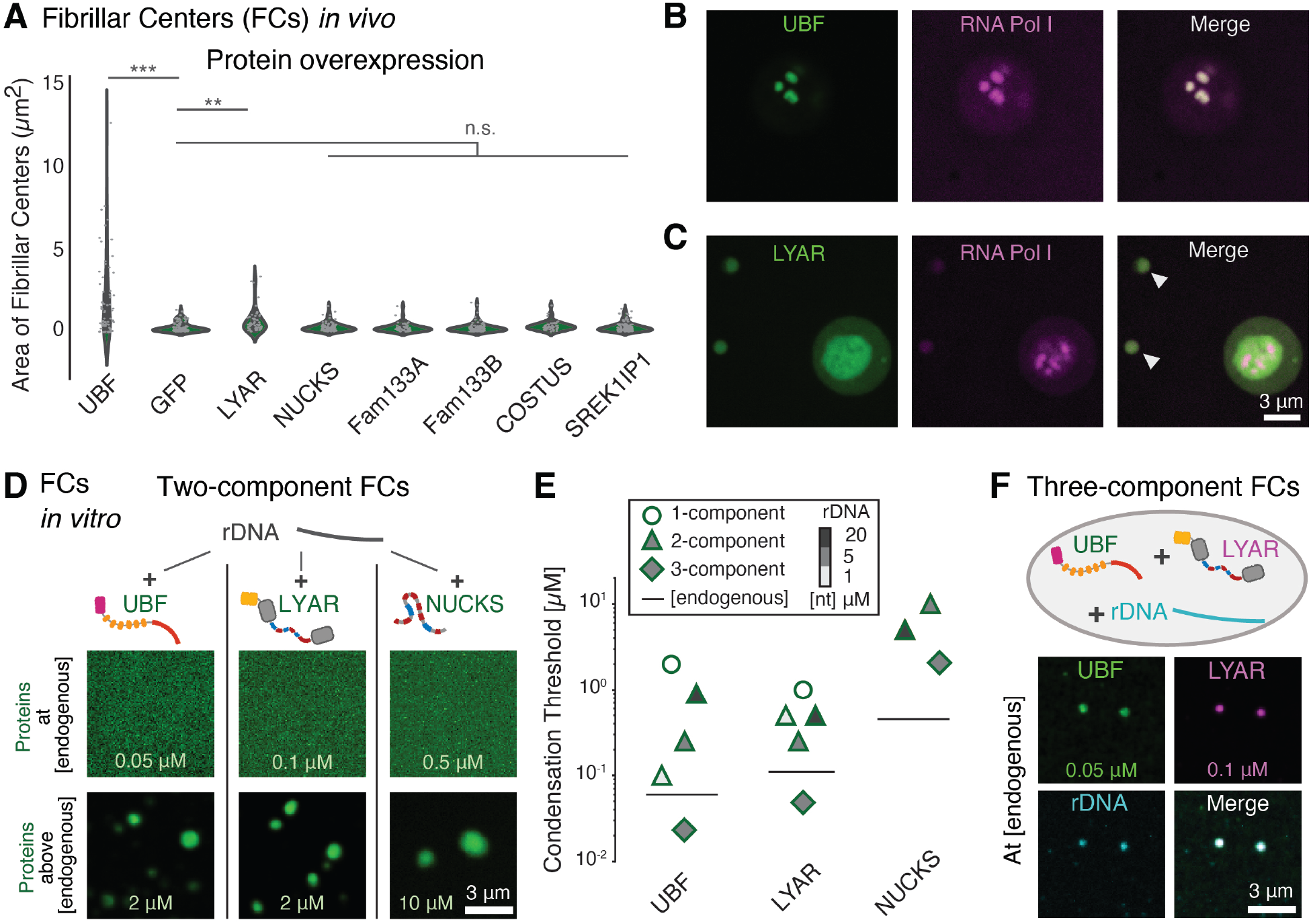
UBF and IDR-containing proteins drive FC condensation. (**A**) Violin plots of FC size from Xenopus oocytes overexpressing GFP-tagged versions of the indicated FC-localized proteins; n.s. denotes p-values < 0.05; ** p-values < 10^−4^, *** p-value < 10^−5^. (**B**) Representative images of oocyte nucleoli overexpressing GFP-tagged UBF and (**C**) GFP-tagged LYAR; Arrowheads indicate *de novo* condensates. (**D**) Two-component mixtures consisting of rDNA (5 μM [nt]) (untagged) and UBF, LYAR, or NUCKS (fluorescently tagged) at the indicated concentrations. **(E)** Threshold concentrations for condensation of recombinant UBF, LYAR, and NUCKS either alone (one-component – unfilled circle), with increasing concentrations of rDNA (two-component – filled triangles), or with three components. The diamonds in columns 1 and 2 show the threshold concentrations of UBF and LYAR that were needed to observe co-condensation in three component mixtures. These thresholds lie below the endogeneous levels of the relevant proteins, shown in each column as a horizontal bar. Condensation was not observed for NUCKS alone or with 1 μM [nt] rDNA. (**F**) Representative condensate formed in three-component mixtures of 5 μM rDNA measured in concentration of nucleotides, UBF (0.05μM), and LYAR (0.1 μM)). The proteins are at their endogenous concentration. Scale bar for all images = 3 μm.

The *in vivo* results point to UBF and / or LYAR being co-scaffolds of the FC along with ribosomal DNA (rDNA). Accordingly, we asked if these components condense and organize into FC-like facsimiles *in vitro*. For these experiments, we used endogenous levels of 5’ETS rDNA. In binary mixtures with rDNA, both full-length UBF and LYAR form co-condensates *in vitro* (Fig. 2D). Despite the presence of six HMG domains in UBF, its condensation with rDNA was not strongly dependent on DNA sequence (Fig SI7). This is consistent with the high affinity of UBF and HMG domains for non-specific DNA ^22^. Nucleolar Ubiquitous Casein Kinase Substrate (NUCKS), a disordered protein consisting entirely of K-blocks+ERRs, also formed condensates with rDNA, suggesting that this grammar and / or the presence of HMG domains is sufficient for co-condensation with rDNA (Fig. 2D).

Does UBF form condensates and recruit rDNA or does it co-condense with rDNA? To answer this question, we quantified the threshold concentration for condensate formation via purely homotypic interactions. For UBF, the threshold concentration for condensation is at least an order magnitude larger than the endogenous levels (Fig. 2E). The same is true for LYAR and NUCKS. This rules out a condensation and recruitment mechanism being operative *in vivo*. Next, we investigated condensates formed in two-component systems. For each of the three concentrations of rDNA that were used, the three systems studied namely, UBF + rDNA, LYAR + rDNA, and NUCKS + rDNA required threshold concentrations of the relevant proteins that are above their endogeneous levels. We then investigated three-component mixtures comprising rDNA, UBF, and LYAR. The measured threshold concentrations of rDNA, UBF, and LYAR that were required to form minimal, FC-like condensates was found to be below the endogeneous levels. Therefore, even in the minimal system comprising three components, an appropriate blend of homotypic and heterotypic interactions is required to generate condensates at concentrations that are prevalent *in vivo*. The intrinsic and co-condensing abilities of proteins *in vitro* follow the hierarchy of UBF > LYAR >> NUCKS. This hierarchy tracks with the ability of the proteins to enlarge FCs when they are overexpressed *in vivo*.

Next, we sought to understand how UBF and proteins such as LYAR and NUCKS with their K-blocks+ERRs drive the assembly of FC-like facsimiles. Using mass photometry and gel electrophoresis, we found that recombinant UBF is mainly a tetramer at endogenous concentrations in physiological buffers (Figs. 3A and SI9). This result is consistent with earlier studies that focused on endogenous or truncated recombinant UBF ^23,24^. Tetramerization is driven by a distinct N-terminal domain that does not involve the HMG domains or the D/E tract. It follows that each UBF tetramer features four D/E-tracts and 24 HMG domains.

**Fig 3:**
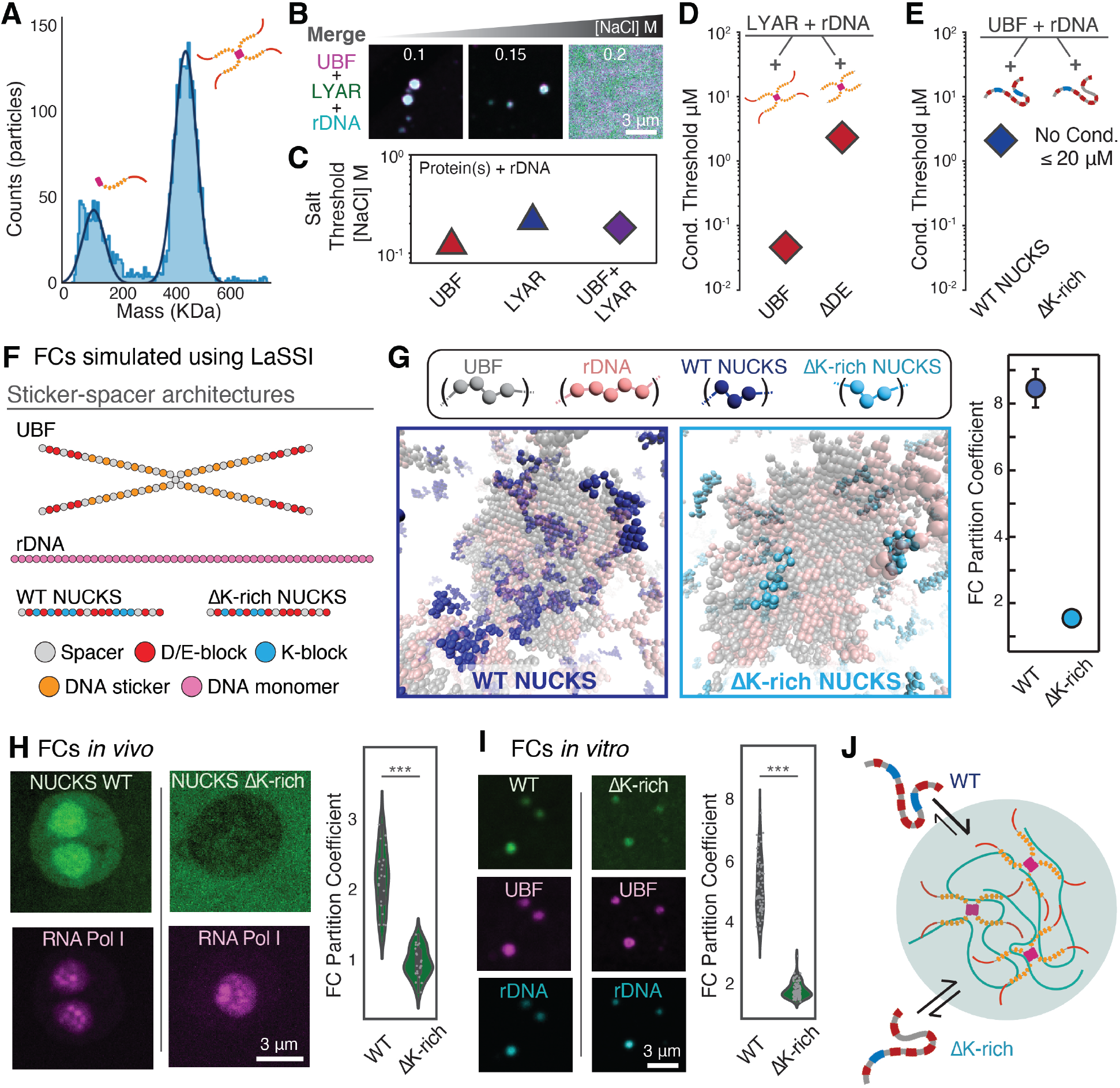
Multivalent K-blocks and D/E-tracts contribute to FC assembly driven by complex coacervation. (**A**) Mass photometry trace of recombinant UBF showing raw counts (light blue histogram) and calculated distribution (dark blue line), which indicates two populations corresponding to the masses of UBF monomers and tetramers. (**B**) Three-component condensates – UBF (0.1 μM), LYAR (0.25 μM), rDNA (5 μM [nt]) – at indicated salt concentrations. (**C**) Threshold salt concentration for condensation of indicated two- and three-component mixtures. All mixtures comprise rDNA. (**D**) Condensation thresholds of WT UBF and ΔDE UBF in three-component condensates with rDNA (5 μM [nt]) and LYAR (0.1 μM). (**E**) Condensation thresholds of WT NUCKS and ΔK-rich NUCKS in three-component condensates with rDNA (5 μM [nt]) and UBF (0.025 μM). (**F**) Coarse-grained architectures of FC components used for LaSSI simulations. (**G**) Representative snapshots of simulated FCs involving UBF, rDNA, and WT NUCKS or ΔK-rich NUCKS. Calculated partition coefficients of WT NUCKS and ΔK-rich NUCKS into simulated FCs. Error bars indicate standard errors from the mean across 10 replicates. (**H**) Representative images of nucleoli from live *Xenopus laevis* oocytes expressing GFP-tagged WT NUCKS or ΔK-rich NUCKS; RFP-PolRIe co-expressed to highlight DFC and FC. Violin plots of NUCKS partition coefficient into DFC/FC phases. (**I**) Representative images of FC facsimiles comprising NUCKS (either WT or ΔK-rich), UBF, and rDNA. Violin plots of NUCKS partition coefficient into FC facsimile. *** denotes *p*-value < 10^−5^. Scale bar for all images = 3 μm. (**J**) Model demonstrating that multivalent K-blocks facilitate partitioning into FCs.

UBF-rDNA interactions, as well as the involvement of proteins such as LYAR, highlight the importance of complementary electrostatic interactions in driving condensation. This is reminiscent of a form of condensation that is known as complex coacervation ^25,26^. A telltale signature of complex coacervation is the ability to dissolve condensates at higher salt concentrations ^26^. Accordingly, we titrated the concentrations of monovalent salts and discovered system-specific thresholds for salt concentrations above which the different FC components do not form condensates (Fig. 3B-C). This points to complex coacervation as the operative mechanism for stabilizing FC-like condensates. However, some aspects of the observed complex coacervation are unconventional, and this pertains to the role of the D/E-tracts in tetrameric UBF. Deletion of the D/E tract shifts the apparent isoelectric point of UBF from ∼ 5.7 to ∼ 9.3. So, we expected that the driving forces for complex coacervation would be enhanced upon deletion of the D/E tract. Surprisingly, removal of the D/E-tracts significantly weakens the driving forces for phase separation in UBF-rDNA mixtures. Specifically, for endogenous levels of rDNA, the threshold concentration of the deletion construct of UBF required to drive condensation is 20-fold higher than the threshold concentration of the full-length UBF (Fig SI10A-C). We also found that the ability of UBF to co-condense into FC facsimiles with LYAR and rDNA is reduced by 50-fold when the D/E-tract is removed (Figs. 3D and SI10B-E).

Overall, our results suggest that complex coacervation driven by UBF involves two distinct contributions from the acid-rich D/E tracts of tetrameric UBF. The D/E tract appears to contribute as a negative regulator in interactions of UBF with rDNA. This observation is consistent with reports suggesting that D/E tracts enable HMG domains to effectively find and bind cognate DNA thus minimizing non-specific interactions with non-cognate sites ^27^. Additionally, the D/E tracts of UBF also appear to enable complementary interactions with lysine-rich motifs in K-blocks+ERRs from proteins such as LYAR and NUCKS.

Next, we assessed the contributions of proteins with K-blocks+ERRs to the complex coacervation with UBF and rDNA. NUCKS lacks any substrate-binding domains and consists almost exclusively of K-blocks+ERRs. It is therefore well-suited for investigating IDR-driven complex coacervation in ternary mixtures of rDNA with proteins featuring K-blocks+ERRs and proteins featuring D/E-tracts. Deletion of the conserved C-terminal, K-rich domain (ΔK-rich NUCKS) compromises the formation of condensates (Fig. 3E). This suggests that the multivalence of K-blocks in NUCKS contributes directly to FC condensation. If true, then the multivalence of K-blocks in proteins with K-blocks+ERRs should also enable partitioning of proteins into facsimiles of FCs ^19^. To test for this possibility, we used a lattice-based method for coarse-grained simulations of equilibrium condensates ^28^. In these simulations, we observed that K-blocks in the K-rich region of NUCKS drive its partitioning into dense phases comprising UBF and rDNA (Fig 3F). We also find that the K-rich region of NUCKS drives its partitioning into FCs *in vivo* and FC facsimiles *in vitro* to roughly the same quantitative degrees (Fig. 3H-I).

The highest scoring protein for K-block enrichment in the proteome is the C-terminal IDR in subunit F of RNA polymerase I (PolR1F). In simulations and *in vitro* measurements, we find that this IDR partitions efficiently into FC facsimiles (Fig SI11). Our findings imply a mechanism whereby proteins with IDRs featuring a multivalence of K-blocks partition efficiently into FC regions of nucleoli (Fig. 3J). Our results suggest that proteins with K-blocks+ERRs contribute as the “molecular glue” to collective interactions that enhance co-condensation of rDNA and UBF.

Next, we sought to determine if and how the different FC proteins contribute to establishing and maintaining the hierarchical organization of the FC and DFC phases of nucleoli. NCL is an abundant, conserved, and essential nucleolar scaffold ^29^. In addition to four folded RNA-binding domains, and a large RG-rich IDR, NCL also features the highest scoring disordered D/E-tract in the human proteome (Fig. 4A). The RG-rich IDR helps confer a high affinity for pre-rRNA^30^. NCL along with the protein fibrillarin, which also contains an RG-rich IDR^7^, has been implicated in driving condensation of the DFC phase of the nucleolus.

**Fig 4:**
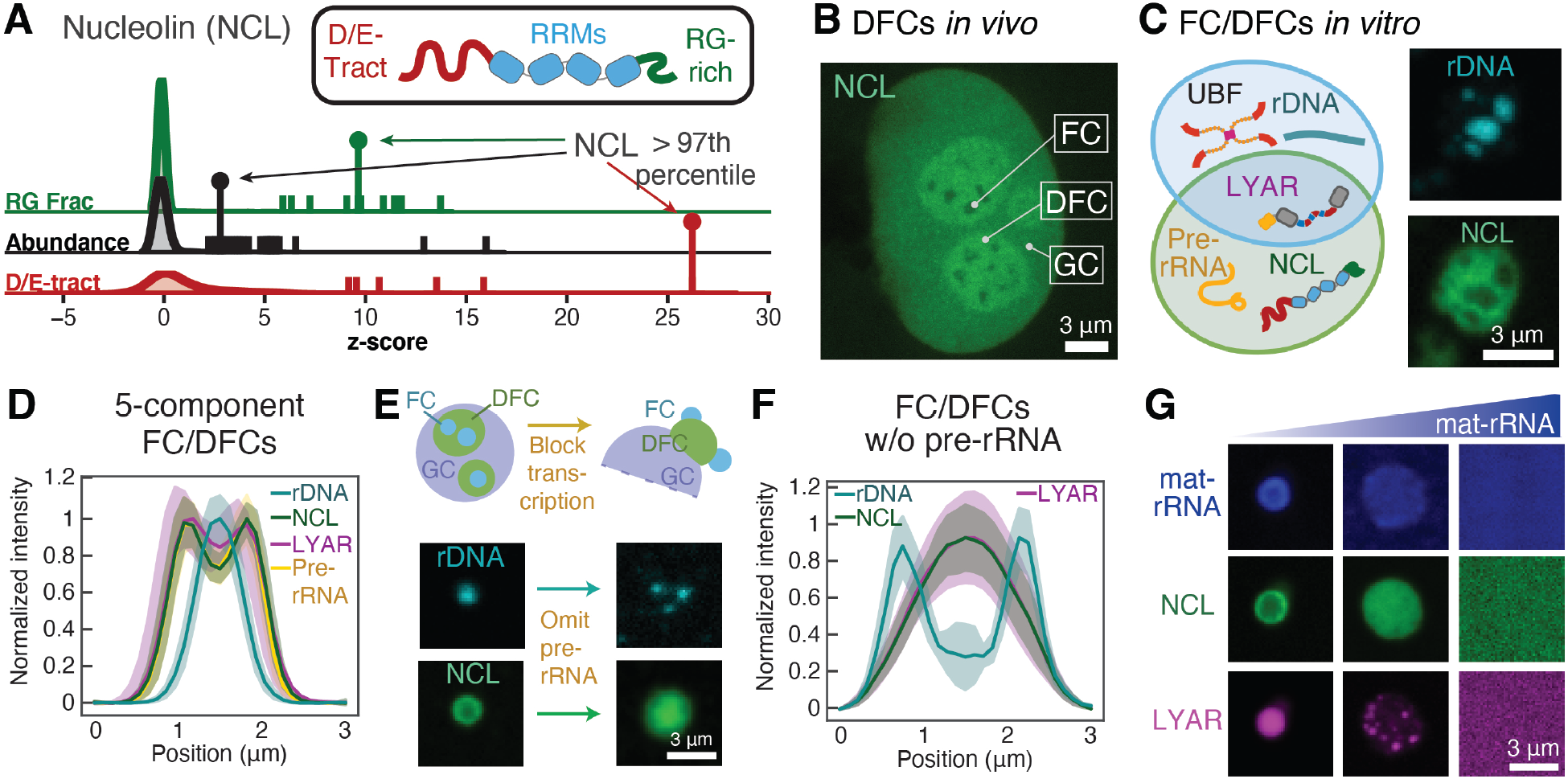
*In vitro* reconstitutions of FCs and DFCs highlight the importance of pre-rRNA-to-mat-rRNA ratios and the contribution of NCL to the FC-DFC interface. (**A**) Schematic of NCL and density distribution graphs showing the relative rank of NCL for fraction of sequence that contains an RG patch among all nucleolar IDRs of length ≥ 30 (7^th^ out of 1951), abundance among all nucleolar proteins (26^th^ out of 781), and D/E-tract (neg-neg patterning) among all nucleolar IDRs of length ≥ 100 (1^st^ out of 433). See methods for additional details. The tall / capped marker indicates NCL; dashes indicate all proteins ranking above NCL and the five closest proteins ranking below it. (**B**) Representative images of nucleoli from a living *Xenopus laevis* oocyte expressing GFP tagged NCL. This highlights the exclusion of NCL from FCs. (**C**) Schematic of the quinary (five-component) mixture and images of multiphase condensates reconstituted *in vitro*. Notice the lack of colocalization between rDNA and NCL. (**D**) Line-scan analysis of condensates formed by the quinary mixture shown in C. (**E**) Schematic of FC extrusion and images of rDNA and NCL in condensates formed by quinary and ternary mixtures. (**F**) Line-scan analysis of condensates in the ternary mixture. In the different line-scan analyses bolded line = median; shaded = 95% confidence interval; n = 25 condensates. (**G**) Images of condensates formed in ternary mixtures containing NCL, LYAR, pre-rRNA (not shown), and mature rRNA at increasing concentrations (1x, 10x, and 30x) of mat-rRNA. Scale bar for all images = 3 μm. Concentrations of UBF and NCL were set at endogenous levels. The concentration of LYAR was 2 μM; see methods for additional details.

Our *in vivo* data show that NCL is excluded from FCs and enriched in DFCs (Fig. 4B). We hypothesized that NCL plays a role in driving the formation of a DFC-like phase that coexists and interfaces with the FC ^7,29^. *In vitro*, we find that, reminiscent of *in vivo* DFCs, recombinant NCL forms well-mixed condensates with pre-rRNA and LYAR (Fig SI12A). The choice of LYAR was motivated by our studies showing that this protein localizes to FCs and DFCs *in vivo*. It has also been implicated as a chaperone for NCL and pre-rRNA^20^. In quinary mixtures comprising UBF, rDNA, LYAR, NCL, and pre-rRNA, with the ratios of the different components set at endogenous levels, we observed a core comprising rDNA and UBF and a shell enriched in NCL and pre-rRNA (Figs. 4C-D and SI12B). This shows that structural facsimiles of coexisting FCs and DFCs can be reconstituted with a minimum of five protein and nucleic acid components.

In addition to essential, multi-functional proteins like UBF and NCL, we find that the establishment and maintenance of facsimiles of FC-DFC interfaces relies on a balance of pre-rRNA and mature rRNA (mat-rRNA) whose levels *in vivo* are controlled by active transcription^4^. Inhibiting pre-rRNA production is known to cause FCs to extrude out into the nucleoplasm whilst remaining partially wetted on a DFC-like region (Fig. 4E) ^4^. Using our *in vitro* facsimiles of coexisting FCs and DFCs, we were able to recapitulate this extrusion phenomenon (Figs. 4E and SI12C/D). All that is required is the exclusion of pre-rRNA. This suggests that the production of pre-rRNA plays a direct role in generating and maintaining coexisting FC and DFC phases.

Lastly, we sought to mimic the transformation of pre-rRNA to mat-rRNA, which is mimicked by adding mat-rRNA at various concentrations, thus changing the mat-rRNA-to-pre-rRNA ratio. Unlike the relatively flexible pre-rRNA, mat-rRNA has a well-defined fold featuring 2’ O-methylation and pseudo uridine modifications ^4^. The two types of rRNA molecules have similar lengths and yet they elicit very different responses in reconstituted systems. We found that mat-rRNA selectively displaces NCL into a region that surrounds the DFC-like phase, thereby generating an incipient flux of mat-rRNA complexed with NCL into the GC^31^. Further, at super-stochiometric ratios of mat-rRNA-to-pre-rRNA, NCL displacement continues until the facsimiles of coexisting FCs and DFCs completely dissolve (Fig. 4G). This reentrant behavior ^32^, driven by mat-rRNA, suggests that in addition to proteins harboring molecular grammars unique to the nucleolus, the relative stoichiometries of pre-rRNA vs. mat-rRNA have a direct impact on the driving forces for assembly of coexisting FCs and DFCs. Our findings are reminiscent of recent observations regarding RNA to DNA stoichiometry in transcriptional condensates ^33^.

Our data suggest that structural facsimiles of coexisting FCs and DFCs can form spontaneously, providing we use the right ratios of pre-rRNA-to-mat-rRNA. This suggests that facsimiles of nucleolar substructures can be set up using networks of interactions among IDRs with D/E-tracts and K-blocks+ERRs, multivalence of substrate-binding domains, and the right ratios of rDNA, pre-rRNA, and mat-rRNA in quinary and higher-order mixtures. As we move towards reconstitutions of functional nucleoli, it will be essential to incorporate transcriptional and rRNA-modifying machinery as well as components such as long non-coding RNAs that help demarcate nucleolar interfaces ^7,34^.

Overall, our work demonstrates the development and deployment of a robust pipeline, driven by a suite of computational approaches that build on improved physicochemical understanding of how IDRs contribute to the specificity of biomolecular interactions ^13^. This approach is likely to help drive reconstitutions of structural facsimiles of other under-investigated cellular condensates. Such reconstitutions will be essential to understand the molecular basis of how different condensates form and coexist while remaining spatially distinct from one another.

## Acknowledgments

We thank Olivia Lazorik and Kaitlyn Hardesty for assistance with cloning and oocyte work. We thank members of the Pappu Lab for insightful feedback and critical reading of the manuscript.

## Author contributions

Conceptualization: MRK and RVP; Methodology: MRK, AZL, KMR, and WO; Investigation: MRK, KMR, MF, and RVP; Visualization: MRK, KMR, and MF; Funding acquisition: RVP, EL; Project administration: RVP; Supervision: MDV, EL, and RVP; Writing – original draft: MRK and RVP; Writing – review & editing: MRK, AZL, KMF, MF, EL, MDV, and RVP.

## Data and materials availability

All data are available in the main text or the supplementary materials.

## Supplementary Materials

Materials and Methods

Figs. S1 to S12

Tables S1 to S4

References (*1*–*15*)

## Supplementary Materials for

## Materials and Methods

### DNA constructs

Detailed information about each DNA construct used in this study can be found in Table S1. DH5α strain *E. coli* cells were used for all subcloning steps and for permanent storage of transfected strains (New England Biolabs (NEB) - C2987I). All DNA constructs were sequence verified by Sanger sequencing (Azenta). Methods for generating DNA and RNA reagents used in reconstitution experiments are described below.

### Protein expression, purification, and fluorescent tagging

All proteins were expressed in Bl21 strain *E. coli* cells (NEB - C2530H) grown either in TB broth (Sigma: T9179) (UBF, NUCKS, and PolR1F IDR) or LB broth (Sigma: L3522) (LYAR and NCL) in Erlenmeyer flasks with ≥ five-fold head volume. Cultures were grown at 37°C at 220 revolutions per minute (RPM) orbital shaking until OD_600_ ∼0.6 was reached. Cultures were then chilled for 15 minutes using an ice bath before being induced with IPTG. Expression-induced cultures in TB broth were grown for 14-18 hours at 18°C; those in LB broth were grown for 6-8 hours at 26°C. Cells were harvested by centrifugation, washed to get rid of residual media, and stored as pellets in 50 mL Falcon tubes at −80°C. Descriptions of the purification procedure of each protein are provided below. Details of the protein sequences – including their amino acid sequence and species origin – can be found in Table S2.

All protein purifications were carried out at 4°C. At the start of a purification a cell pellet was gently resuspended to homogeneity in 35 mL supplemented lysis buffer (protein-specific) then lysed via sonication on a Branson 550 with an L102C horn attachment using five series of the following 20-round cycle: 1 second on / 2 second off at 30% power. Lysis buffer supplements are: 500U DNAaseI (Sigma - 4536282001), 500U RNAse A (Sigma - 10109169001), 5 mg Lysozyme (Sigma - 62971-10G-F), and one protease inhibitor tablet (Sigma - 40694200). After each stage of purification (affinity, ion exchange, and / or size exclusion), protein concentration and nucleotide contamination was assessed on a Nanodrop 2000 via absorbance at 280 nm and measurements of the ratio A260 / A280 of absorbance. Similarly, after each purification stage the inputs, flow-throughs, washes and elutes were examined using SDS-PAGE (sodium dodecyl sulfate– polyacrylamide gel electrophoresis) (Gel: BioRad miniPROTEAN TGX AnyKD - 4569036; MW latter: BioRad Precession Plus Protein Standard Unstained – 1610363), stained with EZblue Coomassie stain (10% (v/v) phosphoric acid, 10% (w/v) Ammonium sulfate, 20% (v/v) Methanol, 1.2% (w/v) Coomassie Blue) and de-stained via serial washes in ddH_2_O.

To express and purify UBF the cells were lysed in supplemented lysis buffer (20 mM Sodium Phosphate, 75 mM NaCl, 20 mM Imidazole, 14.3 mM BME (β-Mercaptoethanol), 200 μM PMSF (phenylmethylsulfonyl fluoride) pH 7.5). Supernatant was recovered from a 25-minute spin at 38,000g and bound to an equilibrated HisTrap FF Crude 5 mL column (Cytiva – 11000458) using an ÄKTA Pure fast protein liquid chromatography (FPLC) module. The NiNTA column was washed in 75 mL lysis buffer. The protein was then eluted in an elution buffer (20 mM Sodium Phosphate, 500 mM NaCl, 350 mM Imidazole, 14-3 mM BME, 200 μM PMSF; pH 7.5). Peak fractions from this affinity purification were pooled and diluted five-fold in dilution buffer (20 mM Sodium Phosphate, 14.3 mM BME; pH 7.5). This solution was further purified via ion exchange chromatography using a contiguous gradient purification protocol with a HiTrap Heparin HP 5mL column (Cytiva – 17040703), Buffer A (20 mM Sodium Phosphate, 100 mM NaCl, 14.3 mM BME; pH 7.5), and Buffer B (20 mM Sodium Phosphate, 100 mM NaCl, 14.3 mM BME; pH 7.5) on the ÄKTA Pure FPLC module. Peak fractions containing His-SUMO-UBF-tev-MBP were pooled and cleaved of SUMO and MBP tags during an overnight dialysis in the presence of 0.02x ULP1 Protease and 0.1x TEV protease in cleavage buffer (20 mM Sodium Phosphate, 300 mM NaCl, 1 mM DTT (Dithiothreitol), pH 7.5). UBF was purified to ≥95% using size exclusion chromatography - HiLoad 16/600 Superdex 200pg column (Cytiva - 28989335) on the ÄKTA Pure FPLC module in storage buffer (22 mM Sodium Phosphate, 1.1 M NaCl, 14.3 mM BME, pH 7.5). The UBF solution was supplemented with 10% glycerol and concentrated in Amicon Ultra 3 MWCO (molecular weight cut-off) concentrator columns (Millipore-Sigma UFC 500396 - hereafter called Amicon concentrators), according to the manufacturer’s suggestions. Concentrated protein was aliquoted into single-use volumes (typically 10 μL), flash frozen in liquid N_2_ and stored at −80°C.

To express and purify LYAR, the Cells were lysed in supplemented lysis buffer (20 mM Sodium Phosphate, 500 mM NaCl, 20 mM Imidazole, 14.3 mM BME, 200 μM PMSF, pH 7). Supernatant was recovered from a 25-minute spin at 38,000g and bound to a 10 mL volume of equilibrated HisPur Ni-NTA resin (Fisher - 88222) in a gravity column. The NiNTA column was washed in lysis buffer until no contaminant protein was detected by Bradford assay. Protein eluted in 20 mL elution buffer (20 mM Sodium Phosphate, 500 mM NaCl, 350mM Imidazole, 14.3 mM BME, 200 μM PMSF; pH 7). Peak fractions from this affinity purification were pooled and diluted five-fold in dilution buffer (20 mM Sodium Phosphate, 14.3 mM BME; pH 7). This solution was further purified via ion exchange chromatography using a contiguous gradient purification protocol with a HiTrap Heparin HP 5mL column (Cytiva – 17040703), Buffer A (20mM Sodium Phosphate, 100mM NaCl, 14.3mM BME; pH 7), and Buffer B (20mM Sodium Phosphate, 100 mM NaCl, 14.3 mM BME; pH 7) on the ÄKTA Pure FPLC module. Peak fractions containing His-LYAR were pooled and purified to ≥95% using size exclusion chromatography - HiLoad 16/600 Superdex 75pg column (Cytiva - 28989333) on the ÄKTA Pure FPLC module in storage buffer (22mM Sodium Phosphate, 1.1 M NaCl, 14.3 mM BME, pH 7.5). The LYAR solution was supplemented with 10% glycerol and concentrated in Amicon concentrators. Concentrated protein was aliquoted into single use volumes (typically 5 μL), flash frozen in liquid N_2_ and stored at −80°C.

To express and purify NUCKS, the cells were lysed in supplemented lysis buffer (20 mM Sodium Phosphate, 500 mM NaCl, 20 mM Imidazole, 14.3 mM BME, 200 μM PMSF, pH 7.5). Supernatant was recovered from a 25-minute spin at 38,000g and bound to an equilibrated HisTrap FF Crude 5 mL column (Cytiva – 11000458) using an ÄKTA Pure fast protein liquid chromatography (FPLC) module. The NiNTA column was washed in 75 mL lysis buffer then protein eluted in elution buffer (20 mM Sodium Phosphate, 300 mM NaCl, 350 mM Imidazole, 14.3 mM BME, 200 μM PMSF; pH 7.5). Peak fractions from this affinity purification were pooled and diluted five-fold in dilution buffer (20 mM Sodium Phosphate, 14.3 mM BME; pH 7.5). This solution was further purified via ion exchange chromatography using a contiguous gradient purification protocol with a HiTrap SP 5 mL column (GE – 17115201), Buffer A (20mM Sodium Phosphate, 100 mM NaCl, 14.3 mM BME; pH 7.5), and Buffer B (20 mM Sodium Phosphate, 100 mM NaCl, 14.3 mM BME; pH 7.5) on the ÄKTA Pure FPLC module. Peak fractions containing His-SUMO-NUCKS-tev-SUMO were pooled and cleaved of both SUMO tags during an overnight dialysis in the presence of 0.02x ULP1 Protease and 0.1x TEV protease in cleavage buffer (20 mM Sodium Phosphate, 300 mM NaCl, 1 mM DTT (Dithiothreitol), pH 7.5). NUCKS was purified to ≥ 95% using size exclusion chromatography - HiLoad 16/600 Superdex 75 pg column on the ÄKTA Pure FPLC module in storage buffer (22 mM Sodium Phosphate, 1.1 M NaCl, 14.3 mM BME, pH 7.5). The NUCKS protein solution was supplemented with 10% glycerol and concentrated in Amicon concentrators. Concentrated protein was aliquoted into single use volumes (typically 5μL), flash frozen in liquid N_2_ and stored at −80°C.

To express and purify the PolR1F IDR, cells were lysed in supplemented lysis buffer (20 mM Sodium Phosphate, 500 mM NaCl, 20 mM Imidazole, 14.3 mM BME, 200 μM PMSF, pH 7). Supernatant was recovered from a 25-minute spin at 38,000g and bound to an equilibrated HisTrap FF Crude 5 mL column (Cytiva – 11000458) using an ÄKTA Pure fast protein liquid chromatography (FPLC) module. The NiNTA column was washed in 75 mL lysis buffer then protein eluted in elution buffer (20 mM Sodium Phosphate, 300mM NaCl, 350mM Imidazole, 14.3 mM BME, 200 μM PMSF; pH 7). Peak fractions from this affinity purification were pooled and diluted five-fold in dilution buffer (20mM Sodium Phosphate, 14.3 mM BME; pH 7). This solution was further purified via ion exchange chromatography using a contiguous gradient purification protocol with a a HiTrap Heparin HP 5mL column (Cytiva – 17040703), Buffer A (20 mM Sodium Phosphate, 100 mM NaCl, 14.3 mM BME; pH 7), and Buffer B (20 mM Sodium Phosphate, 100 mM NaCl, 14.3 mM BME; pH 7) on the ÄKTA Pure FPLC module. Peak fractions containing His-SUMO-PolR1FIDR-tev-GFP were pooled and cleaved SUMO and GFP tags during an overnight dialysis in the presence of 0.02x ULP1 Protease and 0.1x TEV protease in cleavage buffer (20 mM Sodium Phosphate, 300 mM NaCl, 1mM DTT (Dithiothreitol), pH 7). PolR1F IDR was purified to ≥95% using size exclusion chromatography - HiLoad 16/600 Superdex 75 pg column on the ÄKTA Pure FPLC module in storage buffer (22 mM Sodium Phosphate, 1.1 M NaCl, 14.3 mM BME, pH 7.5). The PolR1 F IDR protein solution was supplemented with 10% glycerol and concentrated with Amicon concentrators. Concentrated protein was aliquoted into single use volumes (typically 5 μL), flash frozen in liquid N_2_ and stored at −80°C.

To express and purify the NCL, cells were lysed in supplemented lysis buffer (20 mM Sodium Phosphate, 500 mM NaCl, 20 mM Imidazole, 14.3 mM BME, 200 μM PMSF, pH 7.5). Supernatant was recovered from a 25-minute spin at 38,000g and bound to an equilibrated HisTrap FF Crude 5 mL column (Cytiva – 11000458) using an ÄKTA Pure fast protein liquid chromatography (FPLC) module. The NiNTA column was washed in 75 mL lysis buffer. The protein was then eluted in elution buffer (20 mM Sodium Phosphate, 500 mM NaCl, 350 mM Imidazole, 14.3mM BME, 200 μM PMSF; pH 7.5). Peak fractions from this affinity purification were pooled and diluted five-fold in dilution buffer (20 mM Sodium Phosphate, 14.3 mM BME; pH 7.5). This solution was further purified via ion exchange chromatography using a contiguous gradient purification protocol with a HiTrap Heparin HP 5mL column (Cytiva – 17040703), Buffer A (20 mM Sodium Phosphate, 100 mM NaCl, 14.3 mM BME; pH 7.5), and Buffer B (20 mM Sodium Phosphate, 100 mM NaCl, 14.3 mM BME; pH 7.5) on the ÄKTA Pure FPLC module. Peak fractions containing His-GFP-Tev-NCL (or in some cases His-MBP-Tev-NCL) were pooled and cleaved of GFP during an overnight dialysis in the presence of 0.025x TEV protease in cleavage buffer (20 mM Sodium Phosphate, 100 mM NaCl, 1 mM DTT (Dithiothreitol), pH 7.5). NCL was purified to ≥ 95% using size exclusion chromatography - HiLoad 16/600 Superdex 200pg column (Cytiva - 28989335) on the ÄKTA Pure FPLC module in storage buffer (22 mM Sodium Phosphate, 1.1 M NaCl, 14.3 mM BME, pH 7.5). The NCL solution was supplemented with 10% glycerol and concentrated in Amicon concentrators. Concentrated protein was aliquoted into single use volumes (typically 10 μL), flash frozen in liquid N_2_ and stored at −80°C.

A proportion of purified proteins were covalently conjugated using NHS (N-Hydroxysuccinimide ester) Alexa Fluor™ dye (see Table S3 for specifics on fluorescent dyes used in this study). After the final stage of purification, ∼1 mg of protein was desalted into 100 mM sodium bicarbonate (NaHCO_3_) at pH 8.3 using a PD10 desalting column (GE Healthcare 17085101) according to manufacturer’s suggestions. Desalted protein was concentrated using Amicon concentrators and protein concentration was monitored using 280 nm absorbance on a Nanodrop 2000 (Fisher) periodically. Once protein solutions reached ≥ 5 mg/mL concentrations, they were combined with an NHS-Alexa Fluor™ dye (maintained in DMSO) at a dye-to-protein ratio of 3:1 molarity. This mixture was incubated under gentle rocking for one hour at room temperature in the absence of light. The mixture was then dialyzed overnight into protein storage buffer (20 mM Sodium phosphate, 1M NaCl, 14.3 mM BME, 10% Glycerol at pH 7.5) to remove unincorporated dye. Fluorescently tagged protein was concentrated using Amicon 3 MWCO columns to ≥ 20 μM protein. Concentrations of protein and the covalently attached dye were determined by absorption measurements by nanodrop. Labeling efficiency was calculated from the molar ratios of protein and dye; these values ranged from 0.7-1.2 for all labeled proteins. Labeled proteins were aliquoted into single-use volumes (typically 2 μL), flash frozen in liquid N_2_, and stored at −80°C.

### DNA and RNA reagents

For all DNA reagents, plasmids were prepared using Fisher Pure link quick plasmid miniprep (K210011), according to manufacturer’s instructions. PCR products were purified via PCR clean up (IBI Scientific IB47020) according to the manufacturer’s instructions. To generate tagged DNA, 1% of 405 or 640-labeled dUTP (Table S3) was incorporated into the PCR reaction. Amplified DNA was purified using IBI scientific Gel Clean up (IB47020) according to manufacturer’s suggestions. DNA was eluted in ddH_2_O, and its concentration was measured by absorbance at 260nm on a Nanodrop 2000. It was aliquoted into single use volumes (typically 5μL), flash-frozen in liquid N_2_ and stored at −80°C.

For all RNA reagents other than mat-rRNA, plasmids containing the desired RNA transcript were linearized using restriction enzymes from New England Biolabs according to the manufacturer’s instructions. Linearization was confirmed by gel electrophoresis of DNA samples (control digests where enzyme was omitted were included). Linearized plasmids were purified via PCR clean up (IBI Scientific IB47020) according to the manufacturer’s instructions. RNA was transcribed using the Thermo Fisher mMESSAGE mMACHINE Transcription Kit (using either SP6 polymerase – AM1340 or T7 polymerase – M0255A) according to the manufacturer’s instructions. For all *in vitro* transcription reactions, 100 ng or 1 μg of DNA was used, depending on the length of the *in vitro* transcription (IVT) reaction (100 ng DNA was used for overnight reactions, whereas or 1 μg of DNA was used for 2-4hr reactions). To generate tagged RNA, 1% of Cy3 or Cy5-UTP (Table S3) was incorporated into the IVT reaction. Transcribed RNAs were purified using the NEB Monarch RNA clean up (T2040L), according to the manufacturer’s suggestions. The purity and weight of pure RNA was checked using gel electrophoresis, then aliquoted into single use volumes (typically 2 μL), which were flash frozen in liquid N_2_ and stored at −80°C.

Mat-rRNA is a mixture of rRNAs (5.2s, 18s, and 28s) purified endogenously from xenopus oocyte cytosol via TRIZOL reagent extraction (Fisher - 15596026) according to manufacturer’s instructions. Desiccated RNA pellets were reconstituted in cold ddH_2_O, and their concentration was measured by absorbance at 260 nm using a Nanodrop 2000. From this point on, all procedures took place at 4°C. RNA was either immediately aliquoted into single use volumes (typically 10 μL), flash-frozen in liquid N_2_ and stored at −80°C or used to generate tagged mat-rRNA. For the latter process, extracted RNA of a concentration of at least 7.5 μM was added to freshly prepared sodium periodate and sodium acetate (pH 5). The final concentrations of sodium periodate and sodium acetate are each 100 mM. DEPC-treated ddH_2_O was used to bring the volume of the reaction to 100 μL. The reaction was then incubated for 90 minutes at room temperature in the absence of light. The reaction mixture was then added to an Amicon Ultra-0.5 centrifugal filter with a molecular weight cutoff of 50 kDa, spun for 30 minutes at 14,000 g, and the oxidized RNA was recovered. We then added sodium acetate (pH 5) to a final concentration of 100 mM, and a hydrazide fluorescent dye solution to a final concentration of 1.7 mM. We then added DEPC-treated water to a final volume of 30 μL. The reaction was incubated for 4 hours in the absence of light under gentle agitation. The tagged RNA was then recovered with Amicon Ultra-0.5 centrifugal filters with a cutoff of 50 kDa. The concentration of recovered mat-rRNA was measured by absorbance at 260 nm on a Nanodrop 2000. The material was then aliquoted into single use volumes (typically 2 μL), flash-frozen in liquid N_2_ and stored at −80°C.

### DNA and RNA reagents

For all DNA reagents, plasmids were prepared using Fisher Pure link quick plasmid miniprep (K210011), according to manufacturer’s instructions. PCR products were purified via PCR clean up (IBI Scientific IB47020) according to the manufacturer’s instructions. To generate tagged DNA, 1% of 405 or 640-labeled dUTP (Table S3) was incorporated into the PCR reaction. Amplified DNA was gel purified using IBI scientific Gel Clean up (IB47020) according to manufacturer’s suggestions. DNA was eluted in ddH_2_O, and its concentration was measured by absorbance at 260 nM on a Nanodrop2000. It was aliquoted into single use volumes (typically 5μL), flash-frozen in liquid N_2_ and stored at −80°C.

For all RNA reagents other than mat-rRNA, plasmids containing the desired RNA transcript were linearized using restriction enzymes from New England Biolabs according to the manufacturer’s instructions. Linearization was confirmed by gel electrophoresis of DNA samples (included were control digests where enzyme was omitted). Linearized plasmids were purified via PCR clean up (IBI Scientific IB47020) according to the manufacturer’s instructions. RNA was transcribed using the Thermo Fisher mMESSAGE mMACHINE Transcription Kit (using either SP6 polymerase – AM1340 or T7 polymerase – M0255A) according to the manufacturer’s instructions. For all *in vitro* transcription reactions, 100 ng or 1 μg of DNA was used, depending on the length of the *in vitro* transcription (IVT) reaction (100 ng DNA was used for overnight reactions, whereas or 1 μg of DNA was used for 2-4hr reactions). To generate tagged RNA, 1% of Cy3 or Cy5-UTP (Table S3) was incorporated into the IVT reaction. Transcribed RNAs were purified using the NEB Monarch RNA clean up (T2040L), according to the manufacturer’s suggestions. The purity and weight of pure RNA were checked using gel electrophoresis, then aliquoted into single use volumes (typically 2 μL), which were flash frozen in liquid N_2_ and stored at −80°C.

Mat-rRNA is a mixture of rRNAs (5.2s, 18s, and 28s) purified endogenously from xenopus oocyte cytosol via TRIZOL reagent extraction (Fisher - 15596026) according to manufacturer’s instructions. Desiccated RNA pellets were reconstituted in cold ddH_2_O, and their concentration was measured by absorbance at 260 nM on a Nanodrop2000. From this point on, all procedures took place at 4°C. RNA was either immediately aliquoted into single use volumes (typically 10 μL), flash-frozen in liquid N_2_ and stored at −80°C or used to generate tagged mat-rRNA. For the latter process, extracted RNA of a concentration of at least 7.5 μM was added to freshly prepared sodium periodate and sodium acetate (pH 5). The final concentrations of sodium periodate and sodium acetate were each 100 mM. DEPC-treated ddH_2_O was used to bring the volume of the reaction to 100 μL. The reaction was then incubated for 90 minutes at room temperature in the absence of light. The reaction was then added to an Amicon Ultra-0.5 centrifugal filter with a molecular weight cutoff of 50 kDa, spun for 30 minutes at 14,000g, and recovered the oxidized RNA. We then added sodium acetate (pH 5) to a final concentration of 100 mM, and a hydrazide fluorescent dye solution to a final concentration of 1.7 mM. We then added DEPC-treated water to a final volume of 30 μL. The reaction was incubated for 4 hours in the absence of light under gentle agitation. The tagged RNA was then recovered with Amicon Ultra-0.5 centrifugal filters with a cutoff of 50 kDa. The concentration of recovered mat-rRNA was measured by absorbance at 260 nM on a Nanodrop2000. The material was then aliquoted into single use volumes (typically 2 μL), flash-frozen in liquid N_2_ and stored at −80°C.

### NARDINI plus bioinformatics analysis of the human IDRome

The human proteome was downloaded using Swissprot *Homo sapiens* database (May 2015, 20882 entries)^1^. Each sequence was analyzed using MobiDB to extract the related disorder data^2^. A residue was considered be part of a disordered region if the consensus prediction labeled it as being disordered. All consecutive disordered stretches of length greater than or equal to 100 were extracted leading to 5124 disordered sequences. We refer to this set of sequences as the human IDRome. All IDR sequences were analyzed for 90 sequence features that have been suggested to be important for the function and / or phase separation of disordered regions^3,4^. The sequence features were split into patterning and composition categories. Patterning features were analyzed using a modified version of NARDINI^4^, which stands for Non-random Arrangement of Residues in Disordered Regions Inferred using Numerical Intermixing. Here, 10^3^ scrambled variants were generated per disordered sequence and each distribution was used to extract z-scores for specific binary patterns. Additionally, the sliding window size, *g*, was set to be 5 and 6, and the mean of the results was used. For patterning analysis, residues were grouped as follows: pol ≡ (S, T, N, Q, C, H), hyd ≡ (I, L, M, V), pos ≡ (K, R), neg ≡ (E, D), aro ≡ (F, W, Y), ala ≡ (A), pro ≡ (P), and gly ≡ (G). Z-scores greater than zero imply the disordered sequence is blockier than expected and z-scores less than zero imply the disordered sequence if more well-mixed than expected for the pair of residue types in question.

In addition to identifying non-random binary patterns within IDR, we also extracted 54 compositional features for each IDR sequence. Most features were determined using localCider^5^. These included the fraction of each amino acid type (20 features); the fraction of positive, negative, polar, aliphatic, and aromatic residues (5 features); the ratios of Rs to Ks and Es to Ds (2 features); the fraction of chain expanding (FCE) residues, the fraction of charged residues (FCR), the net charge per residue (NCPR), the fraction of disorder promoting residues, the hydrophobicity, the apparent isoelectric point, and the polyproline-II propensity (7 features). We also included patch features defined as the fraction of sequence made up of patches of a given residue type or RG pair. Here, a patch had to have at least four occurrences of the given residue or two occurrences of RG and could not extend past two interruptions. We examined 20 patch features as no W patch was found in the human IDRome. Then z-scores for each individual sequence in the human IDRome were calculated using the mean and standard deviation of the entire IDRome.

### Extraction of condensate specific enriched sequence features for bioinformatics analysis

Protein sequences from nucleoli, nuclear speckles, and nuclear bodies were extracted from the Human Protein Atlas (HPA)^6^. Cross-referencing with the human IDRome led to 433, 249, and 246 IDRs located in nucleoli, nuclear speckles, and nuclear bodies, respectively. The datasets from Hubstenberger et al., and Jain et al., were used to determine the protein compositions of P-bodies and stress granules ^7,8^. Cross-referencing with the human IDRome led to 58 and 170 IDRs located in P-bodies and stress granules, respectively. To extract sequence features of IDRs that are enriched in specific condensates, the human IDRome was split into the condensate specific IDRome and the remaining IDRome. Then, the distribution of z-scores for a given sequence feature in the two sets were examined and a *p*-value was calculated using the two-sample Kolmogorov-Smirnov test to determine whether the two distributions were identical. If the *p*-value < 0.05, then the signed log_10_(*p*-value) was calculated. Here, the log_10_(*p*-value) was positive if the mean z-score was larger in the condensate IDRome and negative if the mean z-score was smaller in the condensate IDRome. Thus, a positive log_10_(*p*-value) values imply that compositional feature is enriched in the condensate, or the patterning feature is blockier in the condensate.

### Bioinformatic analysis of Nucleolin sequence features

The fraction of the IDR sequence made up of an RG patch, referred to as the RG fraction, was calculated for all IDRs in the human proteome that are of length **≥** 30. Specifically, each patch must have two occurrences of RG and cannot extend beyond two interruptions. Then the mean and standard deviation of the RG fraction across all human IDRs of length **≥** 30 was calculated and used to determine the z-score of all nucleolar IDRs of length **≥** 30. Nucleolin has the 7^th^ highest RG fraction z-score of all 1951 nucleolar IDRs of length **≥** 30. HeLa cell proteome abundance data was extracted from Nagaraj et al^9^. The mean and standard deviation of this dataset was used to determine the abundance z-score values for all nucleolar proteins in this dataset. Nucleolin has the 26^th^ highest z-score of all 781 nucleolar proteins in the Nagaraj et al. dataset. The z-score values for neg-neg patterning were calculated using the modified NARDINI method reported above for all nucleolar IDRs of length **≥** 100. Nucleolin has the highest neg-neg z-score of all 433 nucleolar IDRs and of all 5124 human IDRs.

### Bioinformatics analysis of Human Proteome Atlas (HPA) dataset

UniProt accessions were used to map proteins to subcellular regions and by blocky charge patterning onto the uniform manifold approximation and projection for dimension reduction (UMAP) coordinates of the HPA dataset (April 2019)^10^. For subcellular location, proteins associated with N. speckles, N. bodies, Nucleoli, and N. fibrillar c. where extracted. For the charge patterning features, the UniProt accessions for the top 18 scoring nucleolar IDRs for pos-pos, pos-neg, and neg-neg were collected and then mapped to the UMAP data for visualization of the cellular localization of the proteins containing these IDRs. Visualization was achieved using the ImJoy HPA-UMAP plugin^11^. Additionally, the nucleoli fibrillar center proteins were extracted from HPA and literature scanning to determine how many of the top 18 pos-pos, pos-neg, and neg-neg where localized to the FC/DFC.

### LaSSI simulations

Simulations were performed using LaSSI, a lattice-based Monte Carlo engine^12^. Monte Carlo moves are accepted or rejected based on the Metropolis-Hastings criterion so that the probability of accepting a move is equal to min(1, exp(-βΔ*E*)), where β = 1 / *kT*. Here, *kT* is the simulation temperature and Δ*E* is the change in total system energy associated with the attempted move. Total system energies were calculated using a nearest neighbor model with pairwise interaction parameters described in Table S4. These parameters were chosen to reflect a sticker-spacer architecture wherein the sticker interactions include HMG binding domains with DNA, as well as oppositely charged blocks of residues with each other. In contrast, like charged blocks of residues form very weak interactions with each other, behaving as spacers that contribute a high excluded volume. All other interactions are of a medium strength, typical of spacers that contribute a low excluded volume. All simulations described in this study were performed at *T* = 55 for 4×10^10^ Monte Carlo steps with 10 replicates for each condition. The simulations described in Figure 3F-G involved 250 NUCKS chains, 100 UBF tetramers, and 150 rDNA chains in a cubic lattice with side length 140 lattice units. The simulations described in Fig. S11 involved 385 PolR1F IDR chains, 100 UBF tetramers, and 150 rDNA chains in a cubic lattice with side length 140 lattice units. Chain architectures were determined by coarse-graining IDRs such that each bead in LaSSI is equivalent to ∼10-15 amino acid residues. All simulations were initialized in a smaller cubic lattice to speed up the equilibration into a single condensate. After initialization, the system was allowed to equilibrate fully, as determined by a plateauing of the total system energy. Although this plateauing typically occurs after about 5×10^10^ Monte Carlo steps, simulations were only analyzed after 2×10^10^ Monte Carlo steps to ensure a fully equilibrated system. All simulations analyzed comprised a single large condensate with a coexisting dilute phase. Partition coefficients (PCs) of different chains were calculated as follows: (1) We calculated the radial density of each macromolecular species in the system with reference to the condensate center-of-mass. (2) We fit the following logistic function to the radial density data:

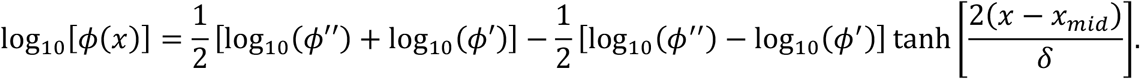

Here, ϕ(*x*) is the calculated radial density, ϕ″ is the density in the dense phase, ϕ′ is the density in the dilute phase, *x*_*mid*_ is the distance between the condensate center-of-mass and the midpoint of the logistic curve, and 5 describes the width of the interface between the dense and dilute phases^13^. (3) Lastly, we calculate the PC as ϕ″/ ϕ′. We perform this routine for all replicates and report the mean value alongside the standard error from the mean for all 10 replicates.

### Image collection and analysis

Confocal imaging was carried out on a Nikon Eclipse Ti2 microscope with a Yokogawa CSU X1 disk module and a LunF laser launch equipped with 405, 488, 561, and 647 lasers. All images shown were taken using a 60X, 1.4 NA Apo oil immersion objective (Nikon) and a Hamamatsu Orca Flash 4.0 CMOS camera. All experiments were carried out at room temperature. NIS-Elements software was used for all image acquisition. All images within a data set were taken with identical imaging parameters ensuring that signal was not saturated (averaging, binning, and projecting were not used). Images were taken and saved as 16-bit ‘.nd2’ files. ImageJ, Python3, and MATLAB were used for all quantitative image analysis (details provided below). All images shown are representative crops of one or a few entities (e.g., a condensate) and the brightness and contrast have been optimized.

### In vivo nucleoli

Harvested oocytes were manually de-flocculated, then subjected to collagenase digestion (gentle rocking) for 2 hours at 18°C. Oocytes were stored in ND96 Buffer (0.005M HEPES, 0.096M NaCl, 0.002M KCl, 0.0018M CaCl_2_, 0.001M MgCl_2_) that was filter sterilized and supplemented with 2.5 mM Sodium Pyruvate C_3_H_3_NaO_3_ (Thermo 11360070) and 1x Penicillin-Streptomycin (Sigma P4333) at 18°C. Healthy stage 6 ocytes were selected and injected using freshly pulled microneedles (Drummond 3-000-203-G/X) and a Drummond Nanoinject II (3-000-204). A total of 23nLs of mRNA in ddH_2_O (typically at a total mass of 20ng) were injected into each oocyte. In all cases, oocytes were injected with mCherry-Pol1Re or GFP-Pol1Re and an mRNA coding for a protein of interest (GFP or RFP tagged). Injected oocytes were stored individually in wells in a 48-well Polystyrene SterileTissue Culture Plates (Fisher FB012930) in supplemented ND-96 buffer at 18°C for at least 18 hours to allow expression and localization of exogenous protein. Immediately prior to imaging, germinal vesicles were manually dissected in mineral oil and mounted on a glass slide with 6 μL of mineral oil. A 22×22mm glass coverslip was gently overlaid onto the sample and nucleoli were then immediately imaged. This procedure was carried out for all proteins of interest for at least two separate harvests of oocytes and similar results were obtained across these biological replicates.

Nucleoli were imaged as a confocal Z-stack. All images shown are of a single Z-slice that is representative and is at least 2 μm above the coverslip; brightness and contrast have been optimized. For more details, see Image collection and analysis section. FC area was calculated as follows. Individual crops of an entire nucleolus were processed using the Otsu method for thresholding in the PolR1e channel (a *bona fide* FC-localizing protein) to create a mask corresponding to the location of FCs. Apparent FCs smaller than 2^2^ pixels (0.10835 μm/pixel) were omitted due to falling below resolution limits. Using the analyze particle and particle manager tools, the FC mask was used to obtain the area (μm^2^) and the mean intensity value (AU) of FCs for both the PolR1e channel and the protein of interest channels. The area obtained using the PolR1e channel was used to report condensate size in the context of expressing various proteins of interest (Figs. 2A, SI6C). The mean intensity values of the protein of interest for individual FCs was saved as ‘FC signal’. To use these values to obtain PCs, we carried out additional analysis as follows: An independent thresholding procedure was applied to duplicated images to obtain the aggregate intensity of all signals that fell below the FC, which we term ‘non-FC signal’. Individual PCs are the difference of the FC signal and non-FC signal, corrected for microscope background. PC values are reported as a normalized value where 1 is the non-FC signal.

For all proteins of interest at least 50 FCs from at least three independent germinal vesicles were analyzed for FC size. Comparative statistics of FC sizes were performed by extracting p-values using 1-tailed Mann-Whitney-Wilcoxon non-parametric test using the MATLAB function mwwtest. Significantly different distributions of sizes were those with *p*-values < 0.05; **denotes *p*-value 10^−4^, ***denotes *p*-value <10^−5^. Similarly, comparative statistics of normalized PC values were performed by extracting *p*-values using 2-tailed Mann-Whitney-Wilcoxon non-parametric test using the same MATLAB function and where ***denotes *p*-value <10^−5^.

### In vitro condensate / phase separation assays

To prepare *in vitro* reconstituted condensates, mixtures of protein(s), DNA, and / or RNA(s) were combined in protein storage buffer (20mM Sodium phosphate, 1M NaCl, 14.3mM BME, 10% Glycerol at pH 7.5) at 10x the final desired concentration. The mixture was then diluted 10x into a no salt buffer (20mM Sodium phosphate 14.3mM BME pH 7.5) to achieve a final [NaCl] of 100mM (roughly physiological conditions). Ten-fold dilutions into higher [NaCl] buffers were used to map the salt condensation thresholds (Fig. 3C-D). All dilutions were performed directly in individual wells in a glass-bottomed 384-well plate (Cellvis-P384-1.5H-N). Immediately upon dilution these mixtures were mixed by pipetting up and down three times. Wells were then sealed, and the contents allowed to equilibrate at room temperature for 20 minutes. After equilibration, the contents of these mixtures were imaged as a confocal Z-stack. All images shown are of a single Z-slice taken at least 1 μm above the coverslip and are representative of the contents of the well; brightness and contrast have been optimized. For more details, see Image collection and analysis section.

PC values of *in vitro* condensates were obtained as follows. For an entire field of view, Otsu thresholding in the channel corresponding to the protein of interest was used to create a mask corresponding to individual condensates. Entities smaller than 5^2^ pixels (0.10835 μm/pixel ≈ ≤0.5 μm in diameter) were omitted. Using the analyze particle and particle manager tools, the mask was used to obtain the mean intensity value (AU) of condensates for all channels. The inverse mask was used to obtain the aggregate intensity of ‘non-condensate’ signal. Individual PCs are the difference of the condensate signal and non-condensate signal, corrected for microscope background. PC values are reported as a normalized value where 1 is the non-condensate signal. Comparative statistics of normalized PC values were performed by extracting p-values using 2-tailed Mann-Whitney-Wilcoxon non-parametric test using the MATLAB function mwwtest. Significantly different distributions of sizes were those with *p*-values < 0.05, ***denotes *p*-value <10^−5^.

### Data visualization

Plots were generated using custom Python 3 scripts; these are freely available upon request. Please email the corresponding author.

### Oligomer analysis of UBF

For both mass photometry and gel electrophoresis analysis of UBF, the protein mixture in storage buffer (20mM Sodium phosphate, 1M NaCl, 14.3mM BME, 10% Glycerol at pH 7.5) at 500 nM [UBF] was diluted 10x into a no salt buffer (20mM Sodium phosphate 14.3 mM BME pH 7.5) to achieve a final [NaCl] of 100 mM and [UBF] of 50 nM (endogenous concentration). This mixture was kept at room temperature for 20 minutes, then analyzed. For gel electrophoresis measurements, equivalent dilutions were also performed into 20mM Sodium phosphate, 1 M NaCl pH 7.5 buffer.

Mass photometry measurements were carried out on a Refyn One Mass Photometer. For a single measurement protein solution (∼10 μL) was applied to a pre-focused slide and counts were recorded for 3 minutes (automated quantification of molecular weight based on airy pattern intensity). The resultant frequency distribution histogram distributions had two peaks at predicted MWs of 109 kDa and 443 kDa, representing monomer and tetramer species of UBF, respectively. For gel electrophoresis measurements, UBF in either 100 mM NaCl or 1 M NaCl was diluted 0.25x into 4x Laemmli SDS sample buffer (Fisher - AAJ60015AC) then immediately run on SDS-PAGE (no sample heating). Gels were stained using Coomassie stain and de-stained. Gels were imaged on a flatbed scanner and processed for optimal brightness and contrast in ImageJ. A line scan of the 100 mM NaCl sample was carried out to determine the positional locations and area-under-the-curve (AoC) values for the monomeric and tetrameric species of UBF. AoC values were obtained by determining the area occupied by each species with the MagicWand tool. Relative amounts of each species were determined using the following: Tetramer relative amount = Tetramer AoC/Tetramer AoC + Monomer AoC; Monomer relative amount = Monomer AoC/Tetramer AoC + Monomer AoC.

### Experimental animals

*Xenopus laevis* (frog) oocytes were used for *in vivo* analysis of protein localization in nucleoli and as source material for biochemical purification of mature rRNA. Mature female *Xenopus laevis* frogs (3-7 years old) are housed in the professional animal facility. Oocyte harvesting is conducted by trained personnel, within the animal preparation room of this animal facility. All procedures are under the supervision of the Washington University Institutional Animal Care and Use Committee Office. All procedures are approved by the Washington University Institutional Animal Care and Use Committee (Animal Welfare Assurance #: A-3381-01). The animal care facility provides additional technical assistance where needed.

**Fig S1.**
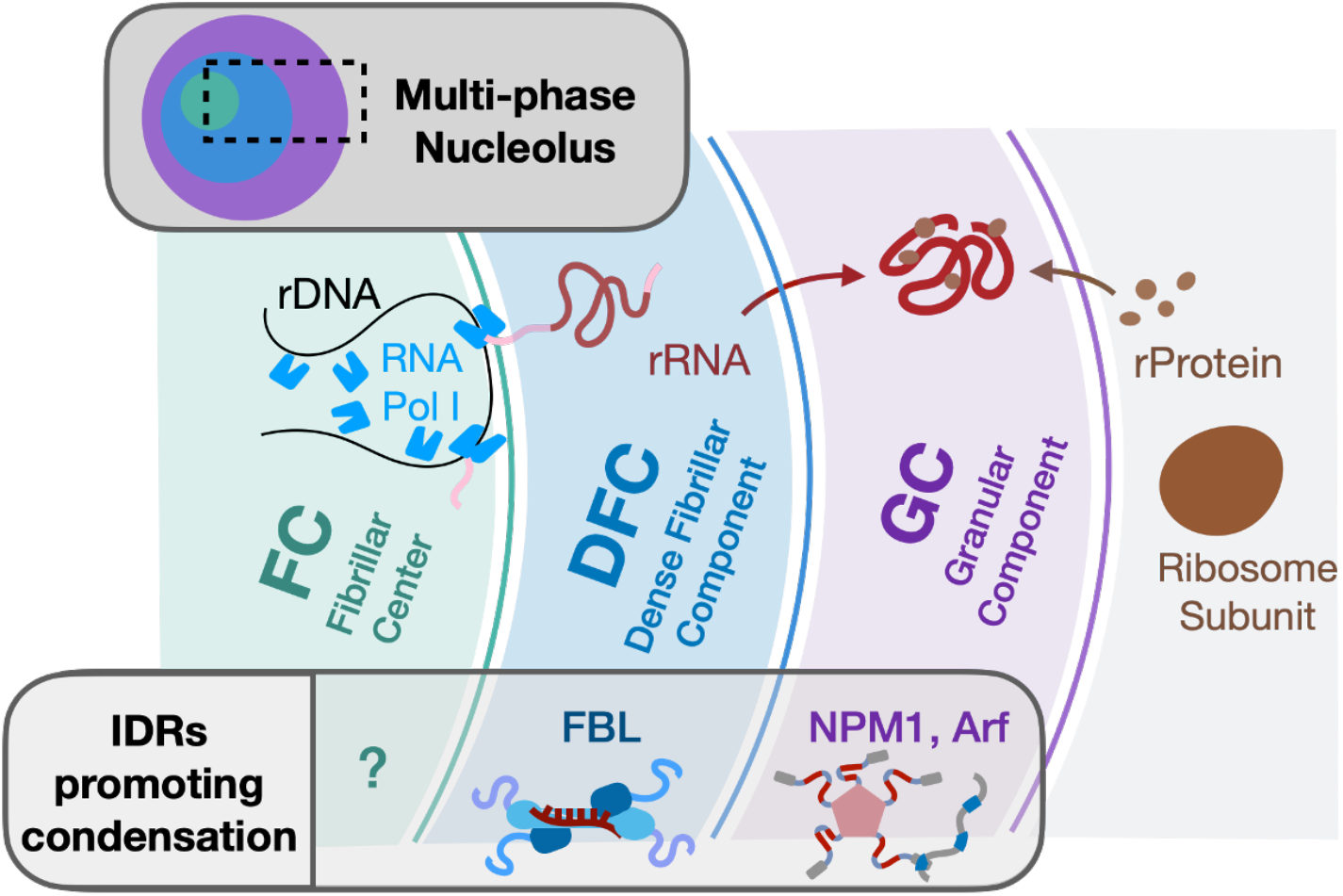
The multiphase nucleolus: Schematic of three-phase nucleoli found in eukaryotes. The inner most phase of the nucleolus, colored teal, is termed the fibrillar center (FC) and is organized around ribosomal DNA (rDNA) loci and known to harbor the transcriptional machinery. Proteins promoting the condensation of this phase have not been proposed or identified in previous studies. Pre-mature rRNA (pre-rRNA), is transcribed at the interface of the FC and the subsequent phase - the dense fibrillar component (DFC), shown in blue. The RG-rich IDR of the FBL subunit of the C/D-box snoRNP (schematized) has been shown to contribute condensation ability *in vitro* and in *in vivo* overexpression assays. rRNA undergoes processing and maturation into a chemically modified and folded mature rRNA (mat-rRNA) in the DFC and the final phase of the nucleolus, the granular component- GC, shown in purple. The outward flux of mat-rRNA into the GC allows it to meet up with the inward flux of ribosomal proteins, and their co-assembly in this phase results in the assembly of proto-ribosomal subunits. The condensation of the GC is thought to involve IDRs on the ribosomal proteins as well as IDRs on NPM1, SURF6 and Arf1 (schematized).

**Fig S2.**
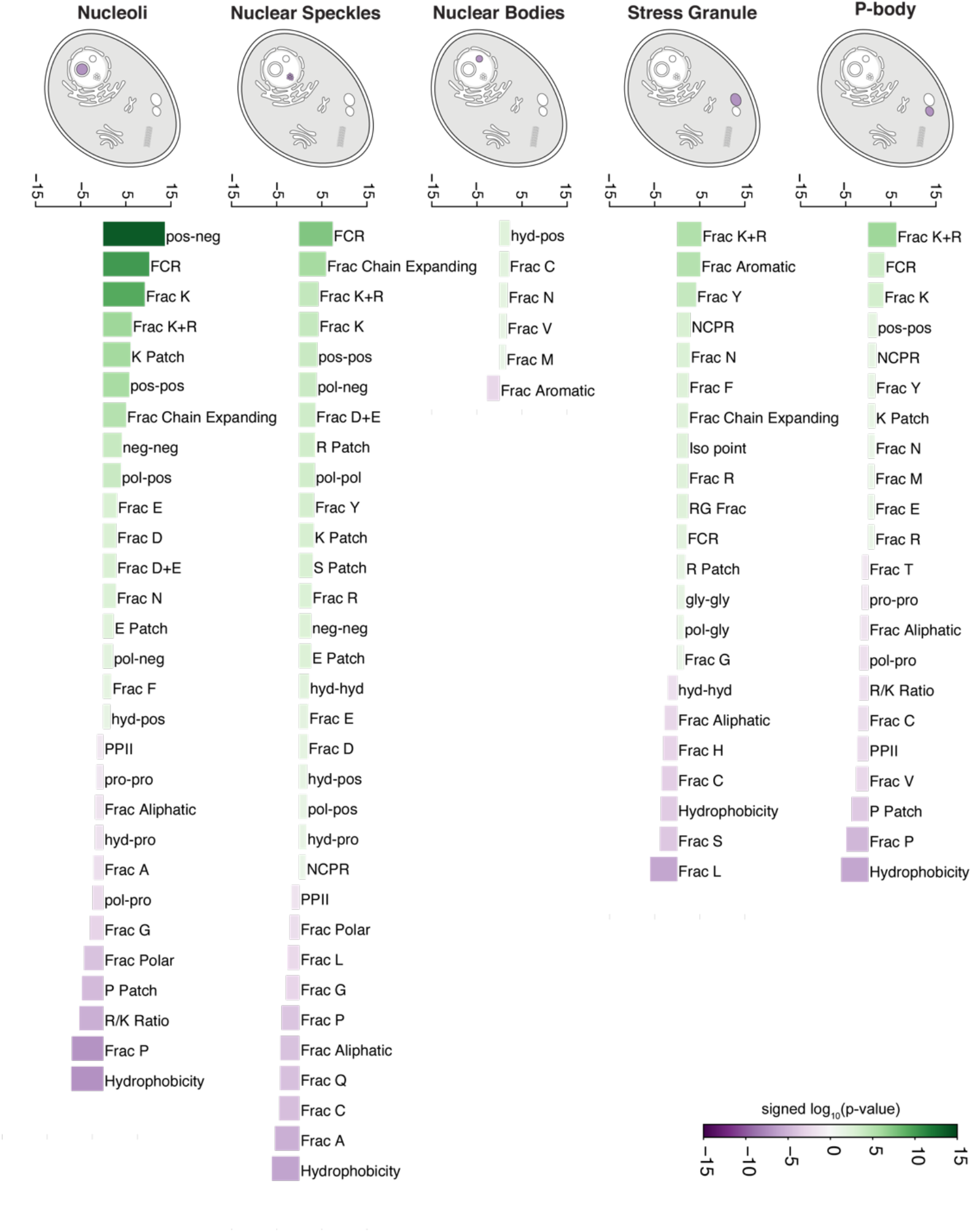
Sequence features of IDRs from proteins drawn from five different biomolecular condensates. Totality of enriched and depleted sequence features within IDRs (length ≥ 100) for proteins drawn from five different condensates (*p*-value < 0.05, two-sample Kolmogorov-Smirnov). Here, the log_10_(*p*-value) is positive if the mean of that feature is higher than the mean of the rest of the IDRome and negative if the mean of that feature is lower than the mean of the rest of the IDRome. Thus, positive values imply that feature is enriched, or the patterning is blockier in the given condensate compared to the rest of the IDRome and negative values imply that feature is depleted, or the patterning is more well-mixed in the given condensate compared to the rest of the IDRome.

**Fig S3.**
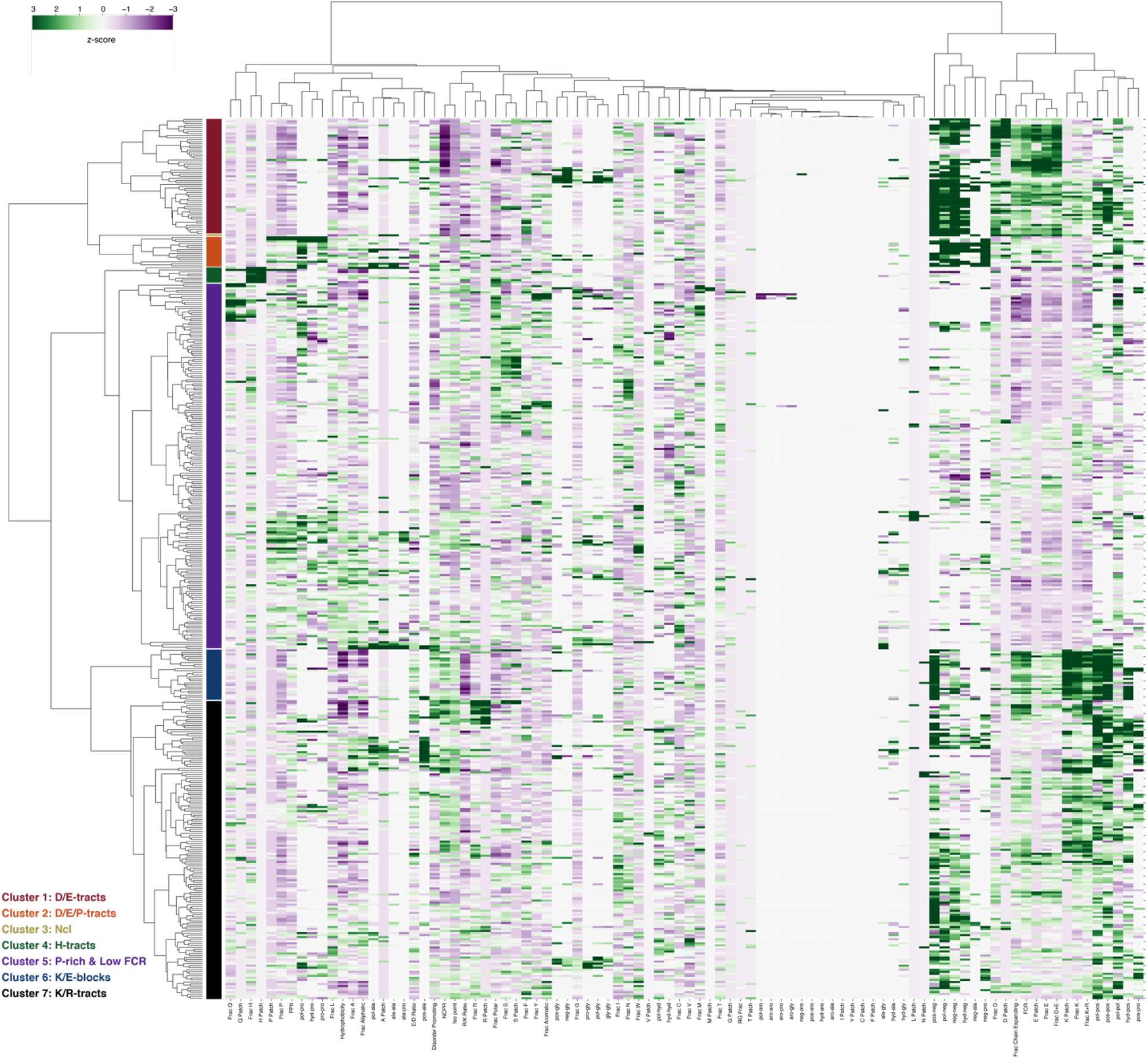
Heat map of all sequence features for all nucleolar IDRs. Sequence feature z-scores for 433 nucleolar IDRs ≥100 amino acids long are hierarchically clustered using the Euclidean distance and Ward’s linkage method. Green colors imply that feature is enriched or blockier than the human IDRome and purple colors imply that feature is depleted or more well-mixed than the human IDRome. Seven clusters were identified: Cluster 1 (red) is enriched in D/E-tracts, Cluster 2 (orange) is enriched in D/E/P-tracts, Cluster 3 (yellow) is Nucleolin, Cluster 4 (green) is enriched in H-tracts, Cluster 5 (purple) is enriched in P and has low FCR, Cluster 6 (blue) is enriched in K/E-blocks, also defined as K-Blocks+ERRs, Cluster 7 (black) is enriched in K/R-tracts. See Fig. S4A for the quantification of sequence feature enrichment for each of the seven clusters.

**Fig S4:**
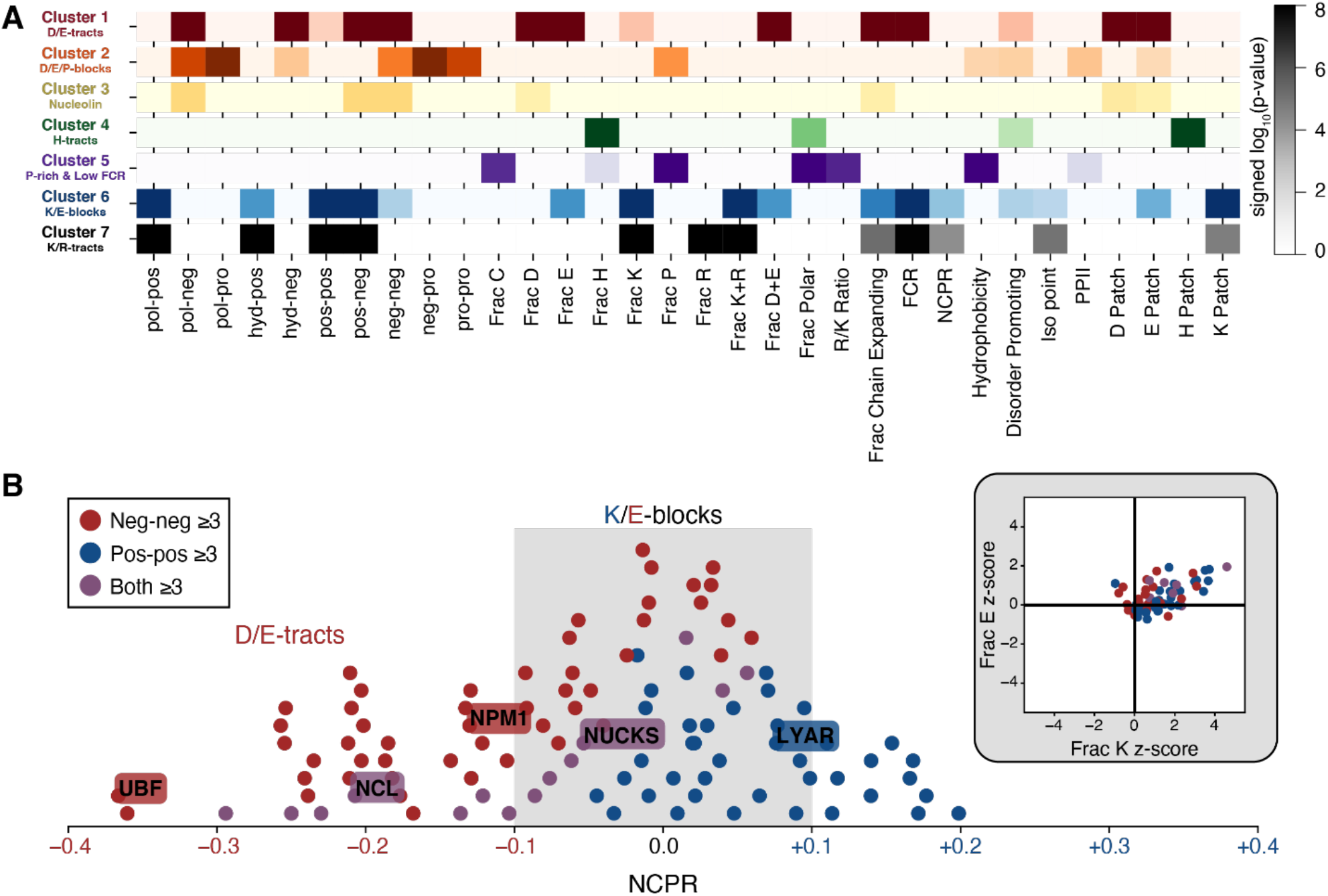
Bioinformatics analyses of predominant Nucleolar IDR sequence features. (A) Reduced heat map of specific enriched sequence features within nucleolar IDRs. Only sequence features with which at least one cluster had a *p*-value < 10^−5^ are plotted. Here, *p*-values are determined by comparing the sequence feature z-score distribution from one cluster to all other nucleolar IDRs using the two-sample Kolmogorov-Smirnov test. Dark colors imply that sequence feature is enriched or blockier in each cluster compared to the rest of the nucleolar IDRome. (B) 104 highest scoring nucleolar IDRs (z-score ≥ 3) for the sequence features neg-neg (red), pos-pos (blue), and both neg-neg and pos-pos (purple) plotted against the net charge per residue of the IDR. Inset shows Frac K z-score versus Frac E z-score for the charge blocky IDRs with |NCPR| ≤ 0.1. Here and in Figure S3, the term K/E-blocks are used interoperable with K-Blocks+ERRs, note that lysines in pos-pos enriched IDRs are blockier, and the same is true for glutamic acids in neg-neg enriched IDRs.

**Fig S5.**
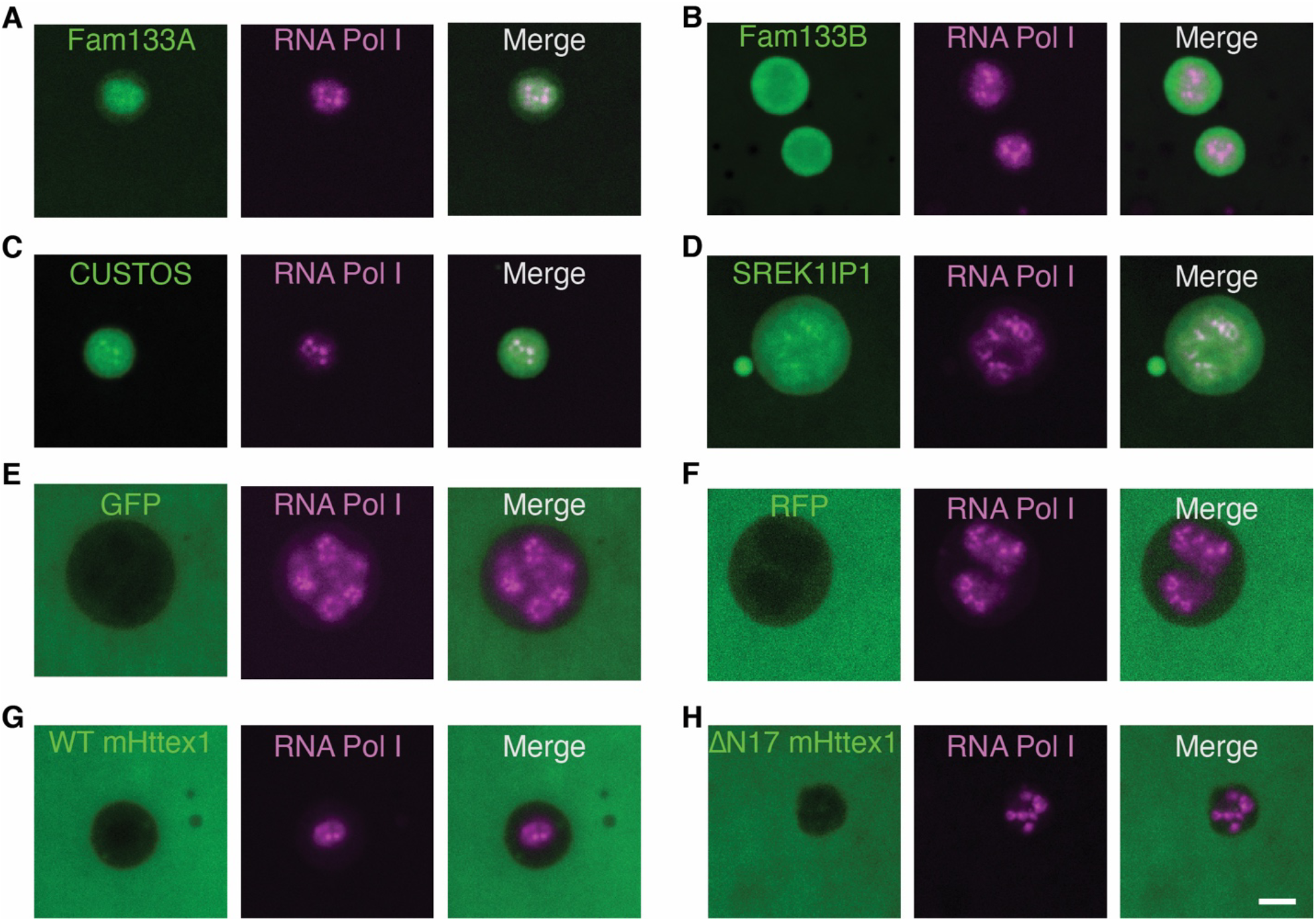
Nucleolar sub-phase localization *in vivo*. Representative images of nucleoli obtained from living *Xenopus laevis* oocytes expressing the indicated FP-tagged protein (**A**) GFP-Fam133A and mCh-PolR1e, (**B**) GFP-Fam133B and mCh-PolR1e, (**C**) GFP-COSTUS and mCh-PolR1e, (**D**) GFP-SREK1IP1 and mCh-PolR1e, (**E**) GFP and mCh-PolR1e, (**F**) RFP and GFP-PolR1e, (**G**) RFP-WT_mHttex1 and GFP-PolR1e, (**H**) RFP-ΔN17_mHttex1 and GFP-PolR1e.

**Fig S6.**
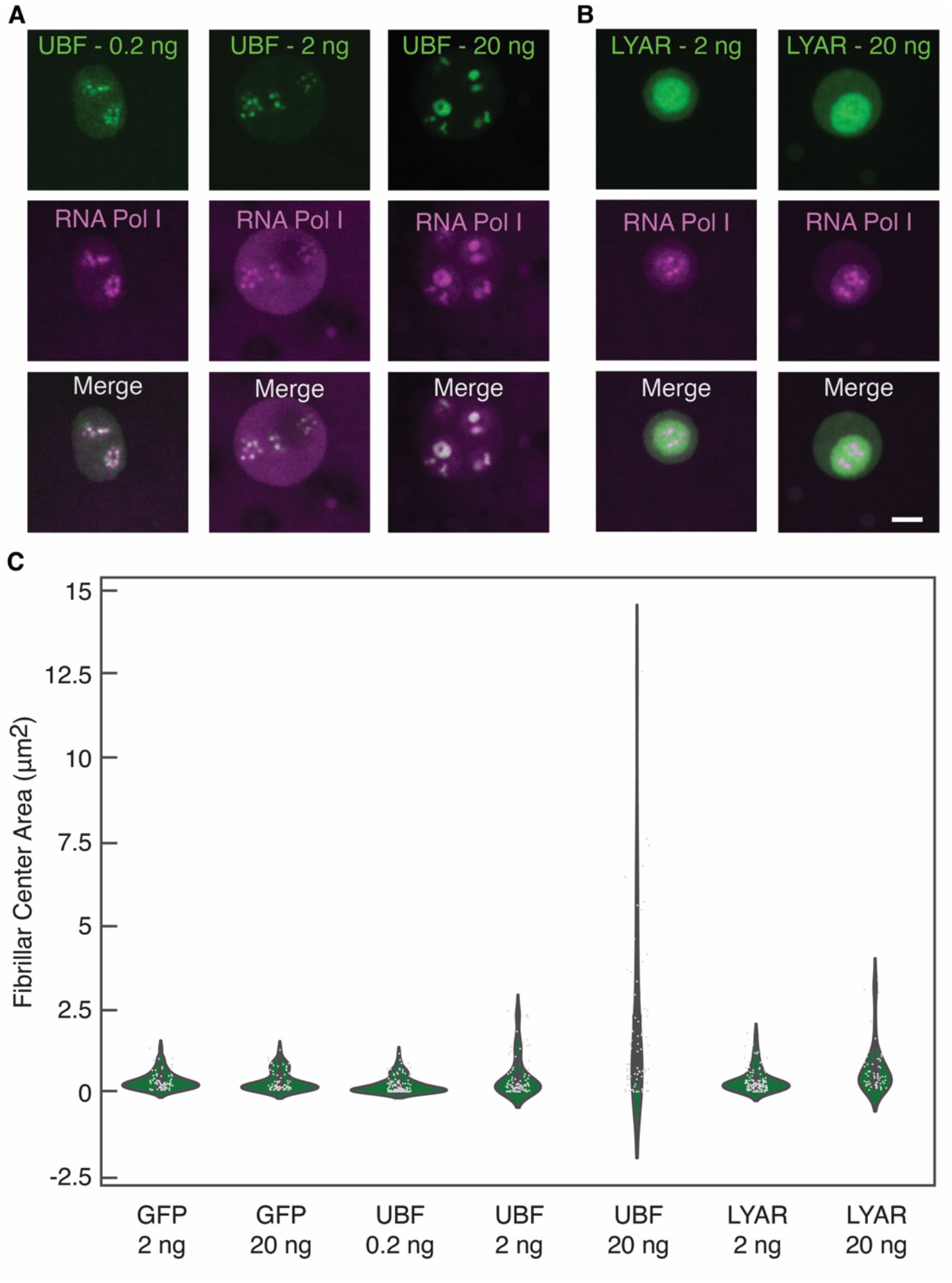
UBF and LYAR over-expression *in vivo*. (**A**) Representative images of nucleoli obtained from living *Xenopus laevis* oocytes expressing GFP-UBF and mCh-RNA_PolI_E. GFP-UBF mRNAs were injected at 0.2ng, 2ng, or 20ng – corresponding columns are indicated. (**B**) Representative images of nucleoli obtained from living *Xenopus laevis* oocytes expressing GFP-LYAR and mCh-RNA_PolI_E. GFP-LYAR mRNAs were injected at 2ng and 20ng – corresponding columns are indicated. In all conditions mCh-RNA_PolI_E mRNA was injected at 2ng. Scale bar = 3 μm. (**C**) Violin plot of Fibrillar Center sizes (area −μm^2^) for indicated overexpression conditions.

**Fig S7.**
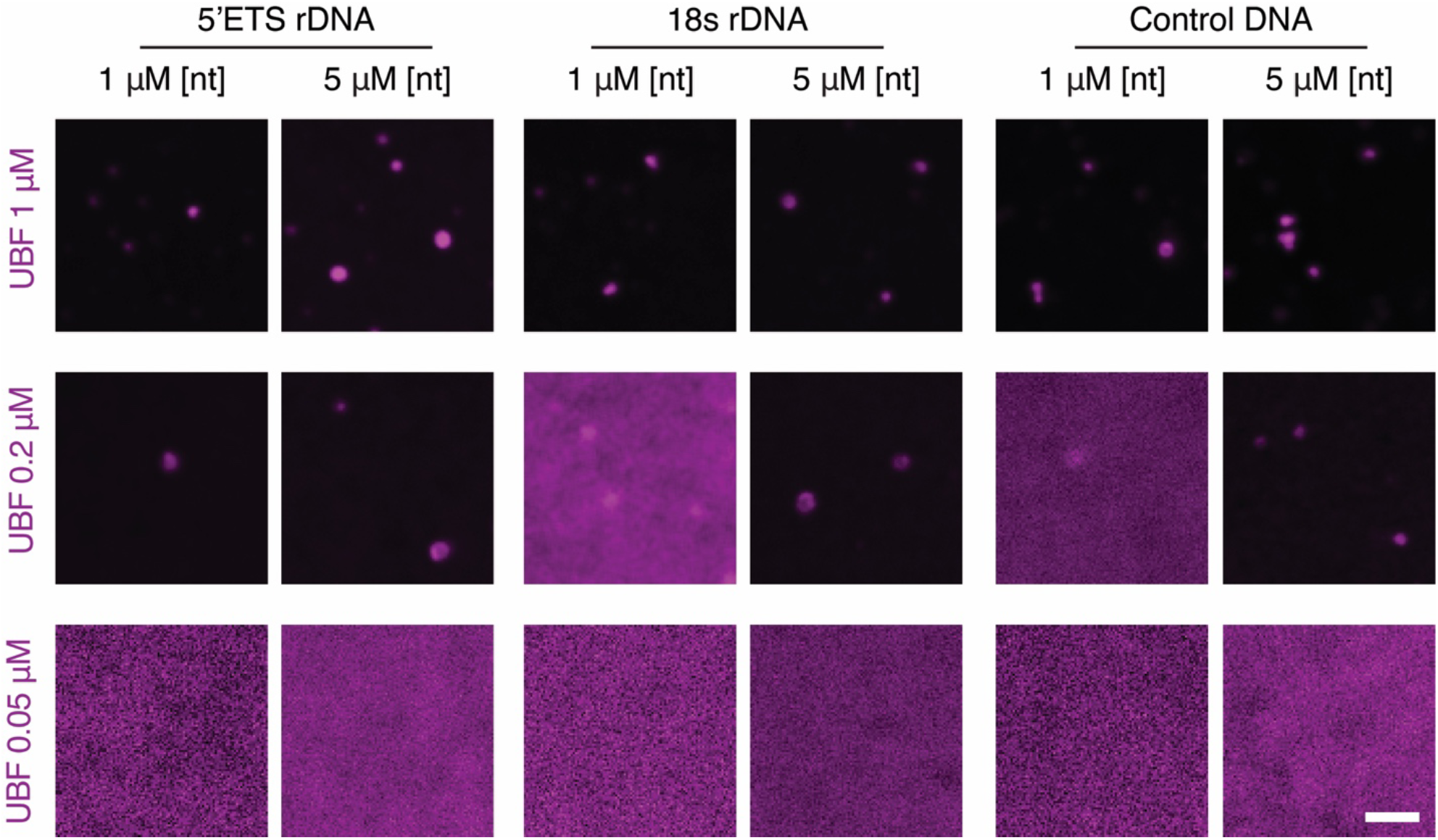
DNA dependence of UBF condensation: Representative images of two-component mixtures of DNA and UBF. The columns show UBF at three different concentrations mixed with: the 5’ETS region of the human rDNA gene (1622bp), the region of the human rDNA gene corresponding to the final mature 18s rRNA fragment (1844bp), and lastly a control DNA fragment corresponding to the Cln3 gene from *Ashbya gossypii* (1612bp), all used at 1 μM or 5 μM [nucleotides]. Scale bar = 3 μm; UBF is N-terminally labeled with Alexa-555.

**Fig S8.**
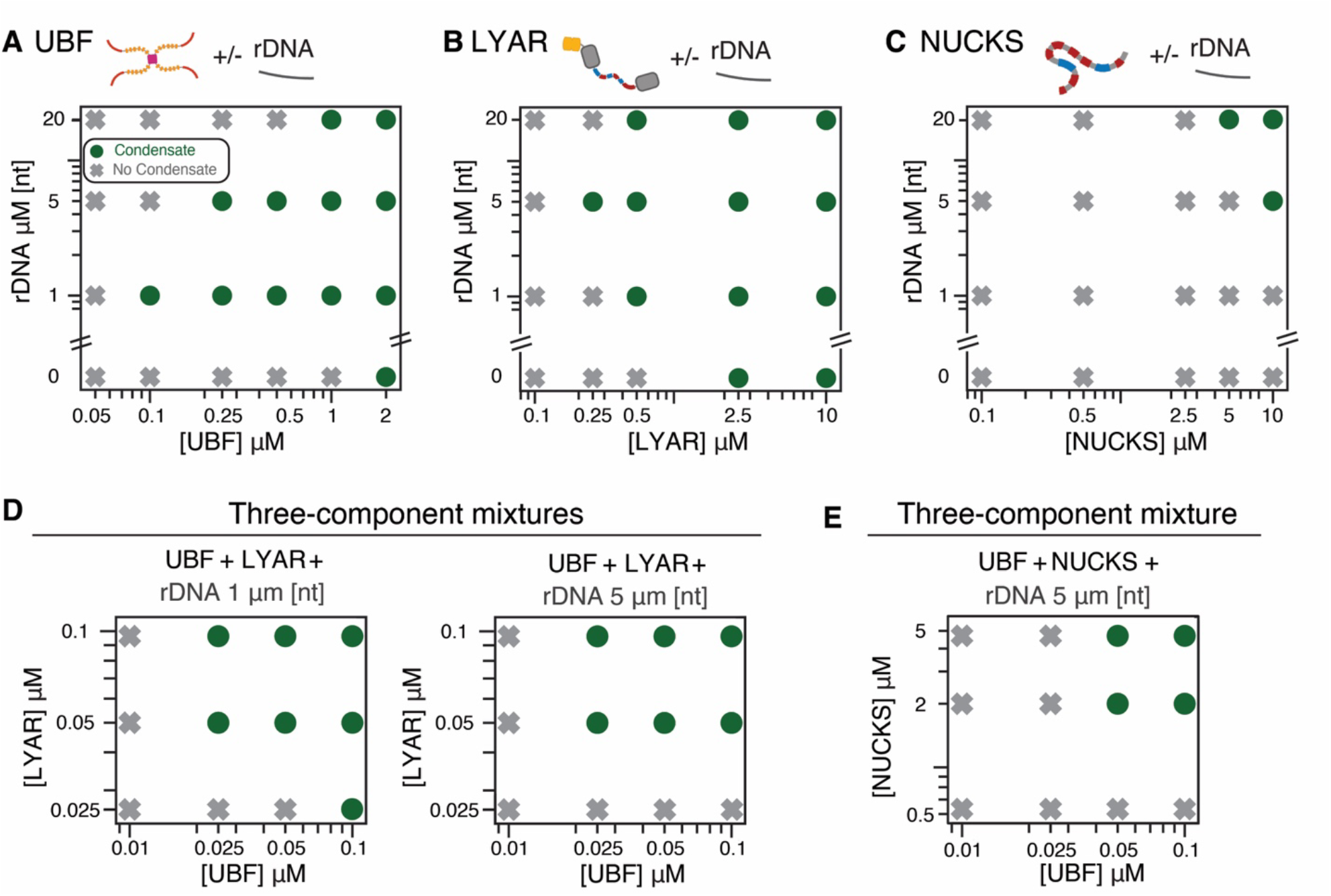
Low concentration arms of the phase boundaries of UBF, LYAR, and NUCKS: Data are shown for (**A**) UBF, (**B**) LYAR, and (**C**) NUCKS as one- and two-component mixtures without or with rDNA at three concentrations – 1, 5, and 20 μM [nt]. (**D**) Low concentration arms of phase boundaries for three-component mixtures of UBF and LYAR mixed with rDNA at 1 μM [nt] or 5 μM [nt]. (**E**) Low concentration arms of phase boundaries of three-component mixture of UBF, NUCKS, and rDNA at nucleotide concentrations of 5 μM.

**Fig S9.**
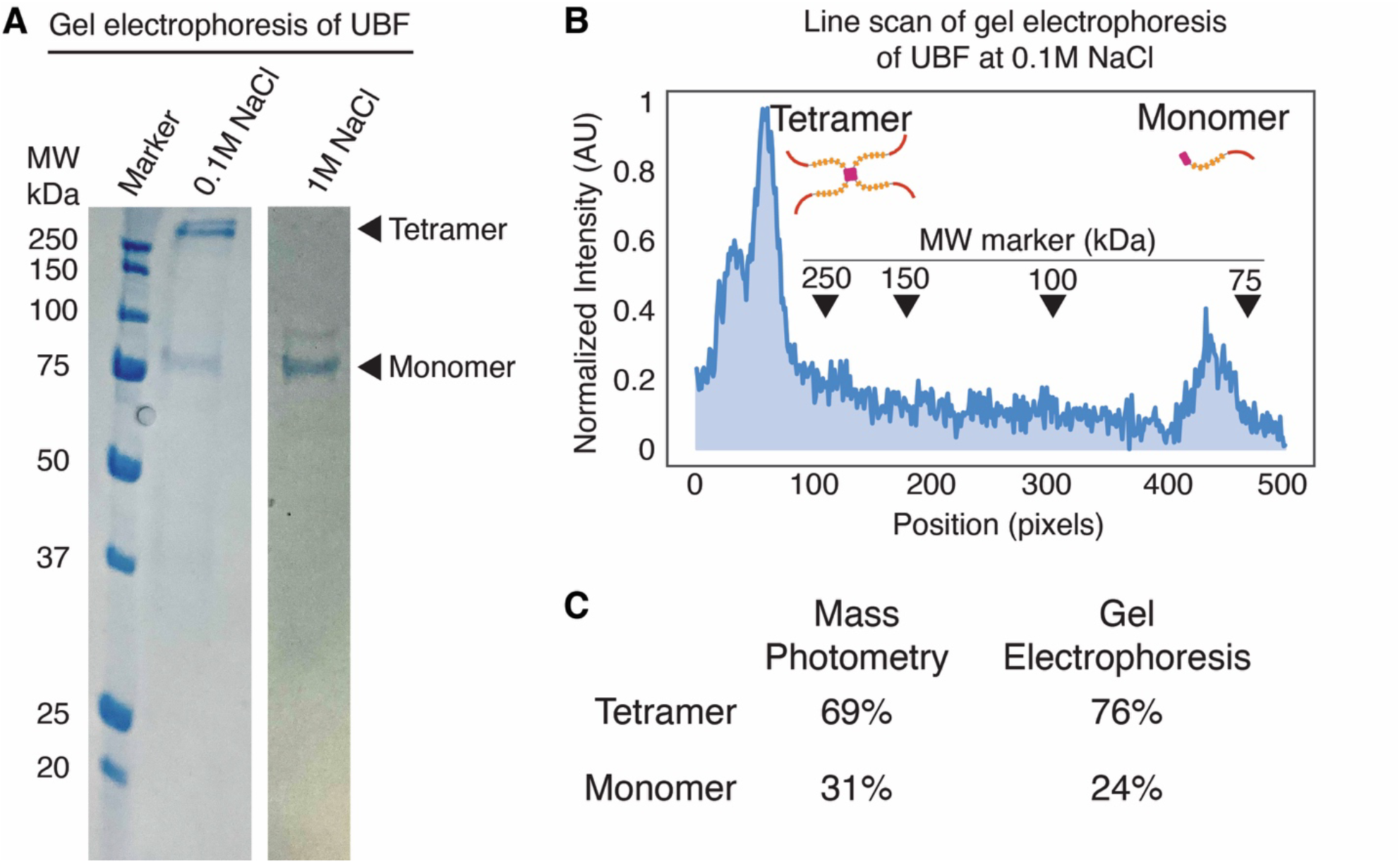
Independent test of UBF tetramer: (**A**) Protein gel electrophoresis of solutions of UBF measured in different concentrations of NaCl. The first solution, shown in lane 2, contains 100 mM NaCl; the second solution, shown in lane 3, contains 1 M NaCl; both contain 20 mM NaPO_4,_ and the pH of the solution is 7.0. (**B**) Densitometry profile of the gel lane containing 100 mM NaCl solution UBF showing protein staining intensity in arbitrary units (AU) normalized to maximum intensity along the gel (position given in pixel units). Arrowheads indicate the position of molecular wight (MW) markers; the expected MW of UBF monomers is 90 kDa and the expected MW of UBF tetramers is 360 kDa. (**C**) Summary of area under the curve calculations for UBF species measured using mass photometry and gel electrophoresis with densitometry.

**Fig S10.**
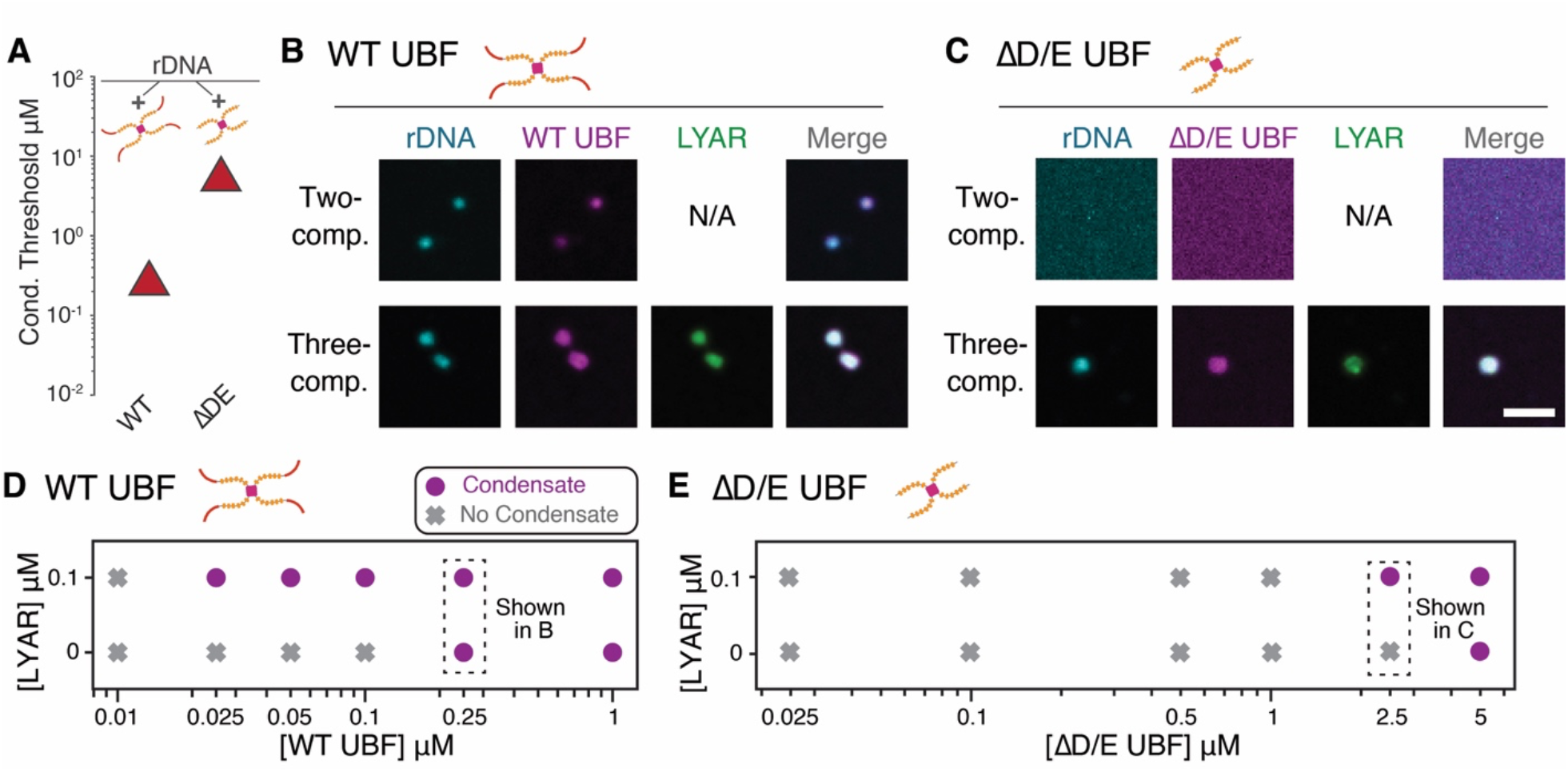
Condensation of UBF WT and UBF ΔD/E, where the D/E tract has been deleted: (**A**) Plotted condensation thresholds of WT UBF and ΔD/E UBF in two component condensates with rDNA (5 μM [nt]). (**B**) Confocal images of mixtures of WT UBF (0.25 μM) in two-component mixture with rDNA (5 μM [nt]) or a three-component mixture with with LYAR (0.1 μM) and rDNA (5 μM [nt]). (**C**) Confocal images of mixtures of D/E-tract UBF (2.5 μM) in two-component mixture with rDNA (5 μM [nt]) or a three-component mixture with with LYAR (0.1 μM) and rDNA (5 μM [nt]); scale bar = 3μm. Phase diagrams of (**D**) WT UBF and (**E**) ΔD/E-tract UBF as a two-component mixture with rDNA (5 μM [nt]) or a three-component mixture with LYAR (0.1 μM) and rDNA (5 μM [nt]).

**Fig S11.**
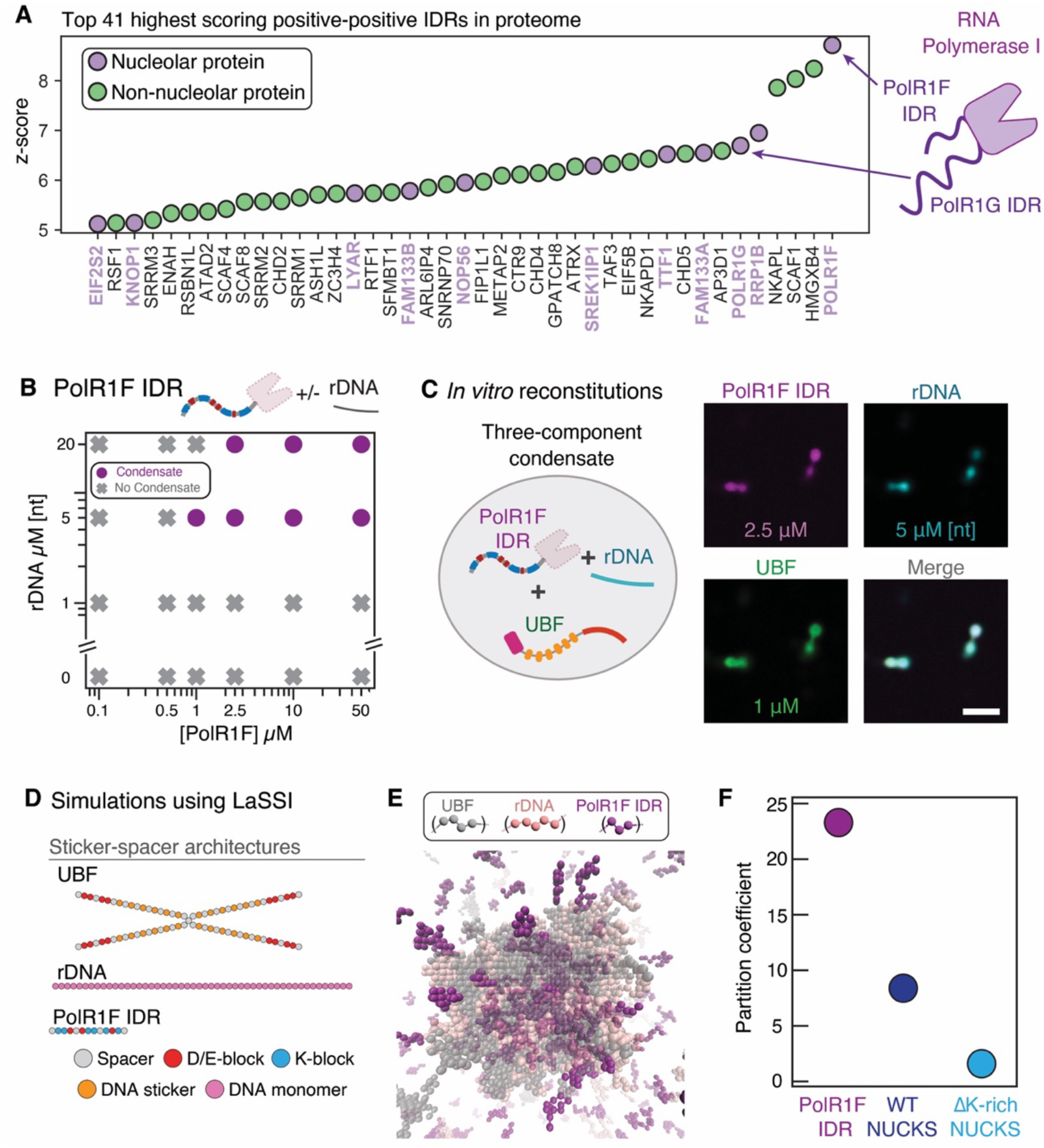
PolR1F IDR is rich in K-blocks and partitions into UBF/rDNA condensates both *in vitro* and in LaSSI simulations: (**A**) Graph of the top 41 highest scoring IDRs in the proteome for the sequence feature positive-positive (z-score >=5). Indicated is the highest scoring IDR in the proteome - the C-terminal IDR of the F subunit of RNA polymerase I (PolR1F) - and the 6^th^ highest scoring IDR in the proteome - the C-terminal IDR of the G subunit of RNA polymerase I (PolR1G). (**B**) Phase diagram of PolR1F IDR in one- and two-component mixtures with rDNA (concentrations indicated) (**C**) Three component mixtures of PolR1F IDR, rDNA and UBF (concentrations indicated); scale bar = 3 μm. (**D**) Coarse-grained architectures of components used for LaSSI simulations (**E**) Representative snapshot of simulated condensate involving UBF, rDNA, and PolR1F IDR. (**F**) Calculated partition coefficients of PolR1F IDR, WT NUCKS, and ΔK-rich NUCKS (the latter two data points are also shown in Figure 3G).

**Fig S12.**
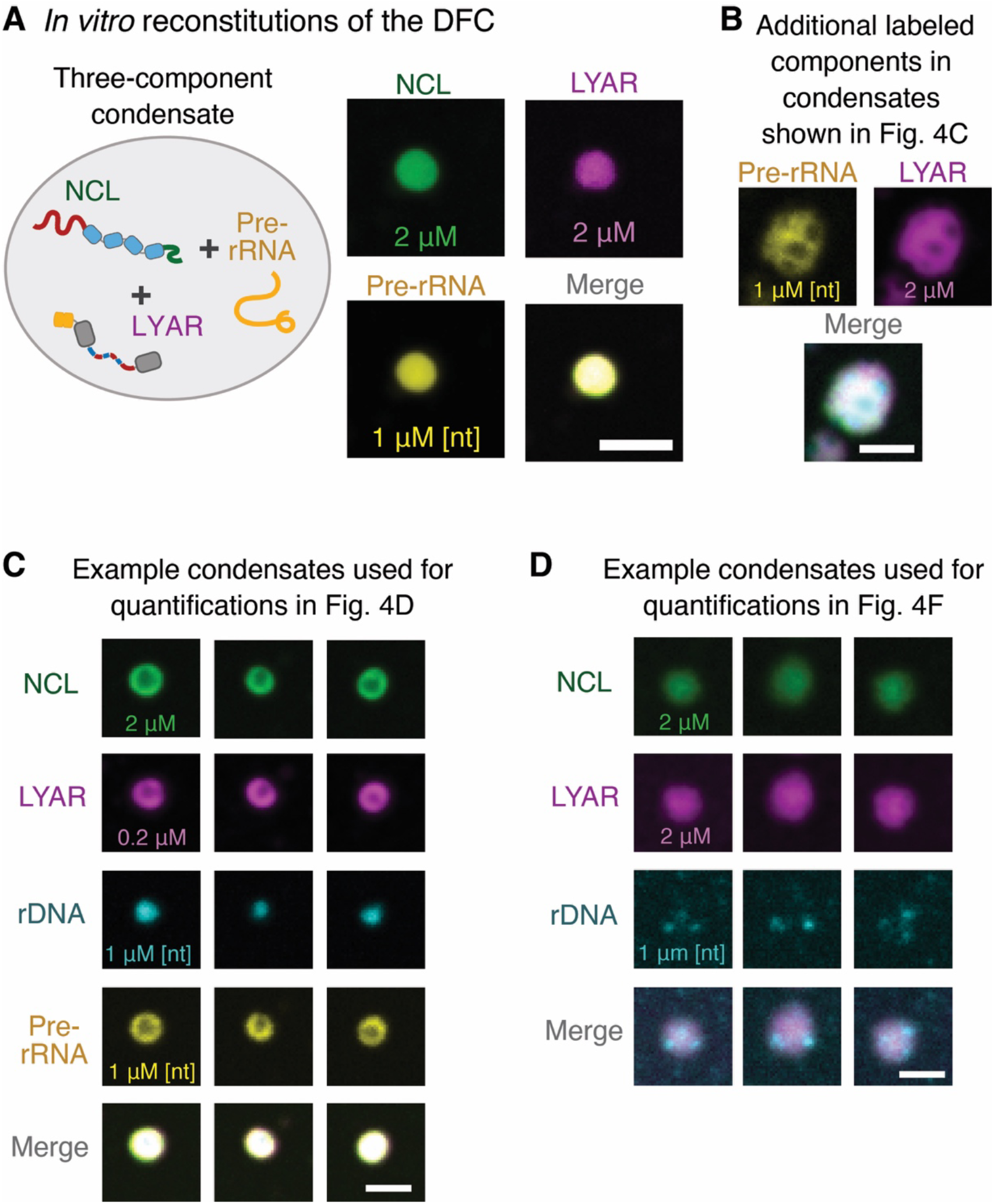
DFC reconstitutions *in vitro*: (**A**) Three component mixtures of NCL, LYAR, and pre-rRNA as minimal *in vitro* reconstitutions of the DFC phase of the nucleolus; concentrations indicated. (**B**) Confocal images of the pre-rRNA and LYAR components of the five-component mixtures displayed in Figure 4C. These mixtures also contain NCL (2 μM), UBF (0.2 μM), and rDNA (1 μM [nt]). (**C**) Confocal images of example five-component FC/DFC condensates used to quantify spatial distribution shown in Figure 4D; [UBF] = 0.2 μM, other concentrations indicated in figure. (**D**) Confocal images of example four-component FC/DFC condensates without pre-rRNA used to quantify spatial distribution shown in Figure 4F; [UBF] = 0.2 μM, other concentrations indicated in figure. All scale bars = 3 μm

**Table S1.** DNA constructs used. List of the DNA constructs used in this study

| Name | Insert description | Vector description | Source | Additional notes |
| --- | --- | --- | --- | --- |
| MK16 | GFP_PolR1e | pCS2+ | Feric et al. 2016<sup>14</sup> | Kind gift from Cliff Brangwynne |
| MK17 | mCherry_PolR1e | pCS2+ |  |  |
| MK20 | NPM1_GFP | pCS2+ |  |  |
| MK21 | NPM1_RFP | pCS2+ |  |  |
| MK39 | His-LYAR (Xenopus) | pET28 | This work | Insert from DNASU - CD00713319 |
| MK49 | GFP-xlNCL (mRNA) | pCS2+ | This work | Insert synthesized to match Xenopus laevis Nucleolin; Uniprot- Q06459 |
| MK50 | mCherry-xlNCL (mRNA) | pCS2+ | This work |  |
| MK51 | GFP_LYAR (mRNA) | pCS2+ | This work | Insert from DNASU - CD00713319 |
| MK52 | mCherry-LYAR (mRNA) | pCS2+ | This work |  |
| MK55 | His-GFP-Tev-NCL | pETDuet-1 | This work | Insert synthesized to match Xenopus laevis Nucleolin; Uniprot- Q06459 |
| MK78 | His-MBP-Tev-NCL | pET28 | This work |  |
| MK89 | SP6_5'ETS_rDNA | pCR-TOPO | This work | Insert derived from PCR amplification of a 1622bp region corresponding to the annotated 5' ETS region of the Human rDNA loci (Gene ID: 6052); This construct was used to generate rDNA reagents via PCR amplification |
| MK90 | SP6_18s_rDNA | pCR-TOPO | This work | Insert derived from PCR amplification of a 1844bp region corresponding to the annotated 18s region of the Human rDNA loci (Gene ID: 6052); This construct was used to generate rDNA reagents via PCR amplification and pre-rRNA reagents using in vitro transcription |
| MK93 | GFP_FAM133A (mRNA) | pCS2+ | This work | Insert from DNASU - CD00353663 |
| MK94 | GFP_FAM133B (mRNA) | pCS2+ | This work | Insert synthesized to match Uniprot Q5BKY9 |
| MK95 | GFP_C12orf43 (mRNA) | pCS2+ | This work | Insert from DNASU - CD00440055 |
| MK97 | GFP_NUCKS (mRNA) | pCS2+ | This work | Insert from DNASU - CD00709868 |
| MK98 | GFP_SREK1IP1 (mRNA) | pCS2+ | This work | Insert from DNASU - CD00714345 |
| MK106 | His-SUMO-NUCKS-Tev-SUMO | pET28 | This work | Insert synthesized to match Uniprot Q9H1E3 |
| MK117 | GFP_NUCKS_ΔKrich (mRNA) | pCS2+ | This work | Cloned from pMK97 via site directed mutagenesis |
| MK137 | His-SUMO-UBF-Tev-MBP | pET28 | This work | Insert synthesized to match Uniprot P17480 |
| MK140 | His-SUMO-PolR1FIDR-Tev-GFP | pET28 | This work | Insert synthesized to match Uniprot Q3B726 |
| MK141 | His-SUMO-NUCKSΔKRich-Tev-SUMO | pET28 | This work | Cloned from pMK106 via site directed mutagenesis |
| pMK152 | His-SUMO-UBFΔDE-Tev-MBP | pET28 | This work | Cloned from pMK137 via site directed mutagenesis |
| pUC19_T7_Cln3 | T7_Cln3 | pUC19 | This work | Insert from Zhang et al 2015<sup>15</sup> (Kind Gift from Amy Gladfelter); DNA used in Figure SI7; a 1612 bp amplicon of Cln3 DNA was generated via PCR amplification |

**Table S2.**
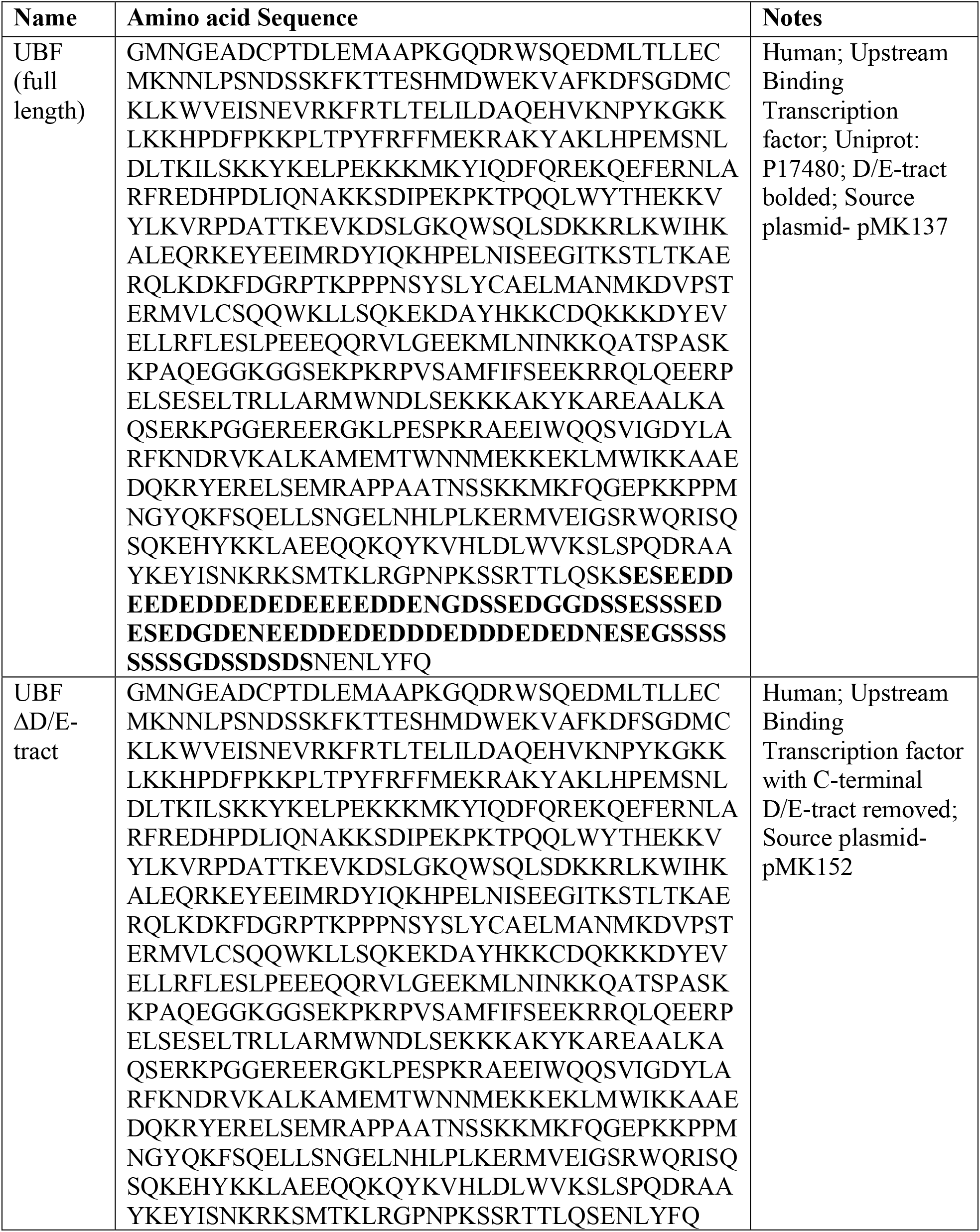
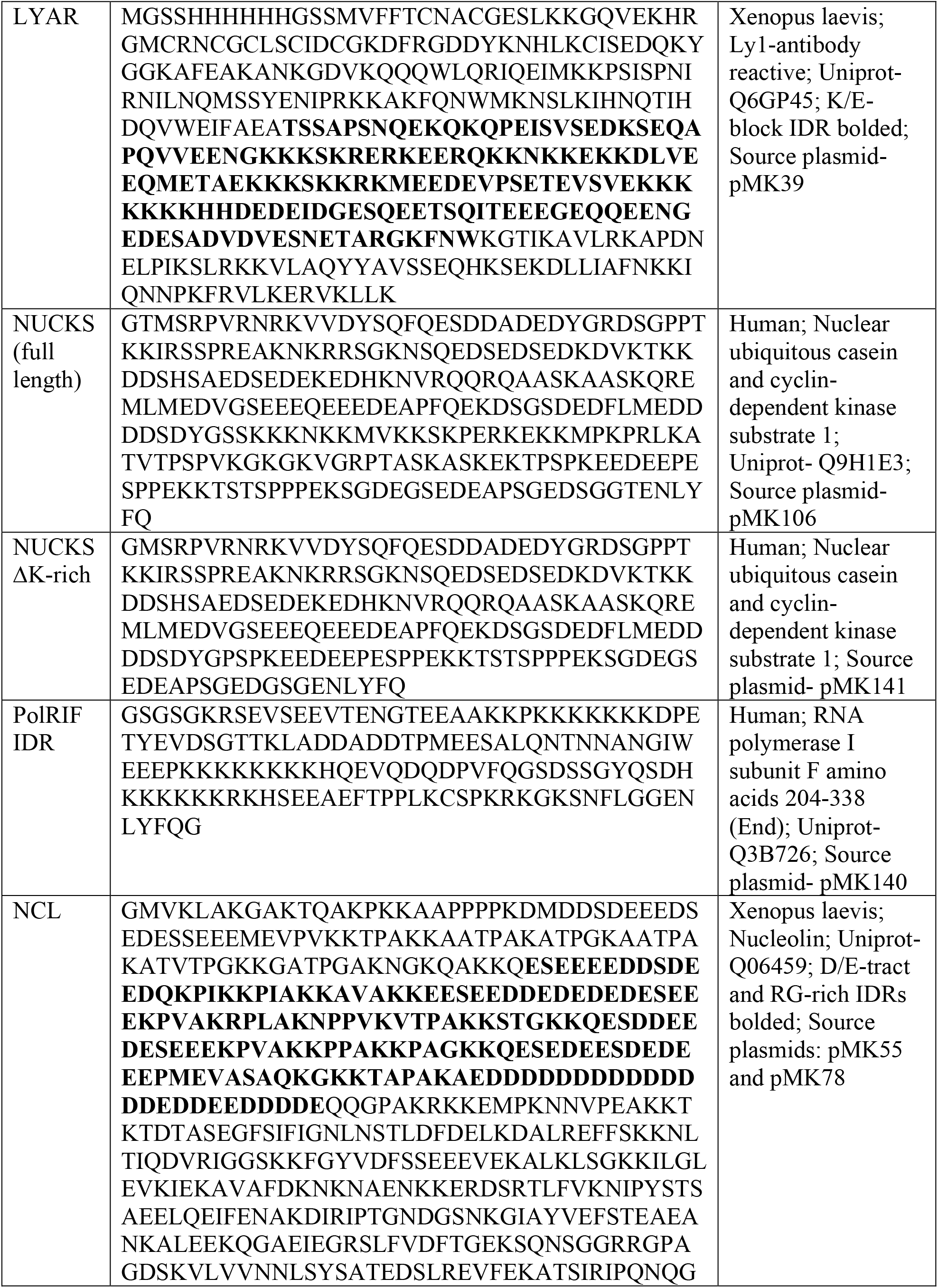
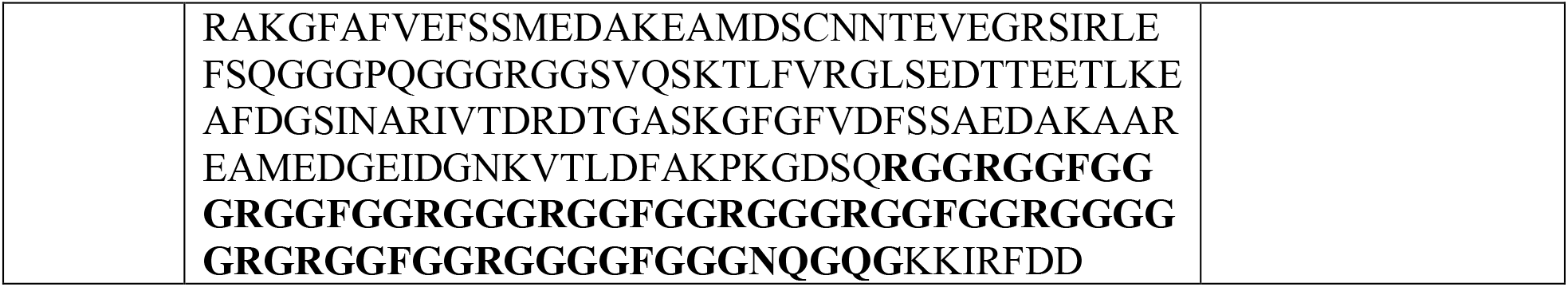
Proteins used. List of the proteins used in this study

**Table S3.** Fluorescent dyes used. Listed are the fluorescent dyes used that were conjugated to proteins or incorporated into RNA and DNA reagents

| **Dye-Conjugate** | **Use** | **Source** |
| --- | --- | --- |
| CF®405M dUTP | Labeling DNA reagents | Biotium - 40100-T |
| CF®640R dUTP | Labeling DNA reagents | Biotium - 40007-T |
| Cy®3-UTP Cytiva | Labeling RNA reagents | Sigma - GEPA53026 |
| Cy®5-UTP Cytiva | Labeling RNA reagents | Sigma - GEPA55026 |
| AlexaFluor™405 NHS Ester | Labeling proteins | Fisher – A30000 |
| AlexaFluor™488 NHS Ester | Labeling proteins | Fisher – A20100 |
| AlexaFluor™555 NHS Ester | Labeling proteins | Fisher – A20009 |
| AlexaFluor™647 NHS Ester | Labeling proteins | Fisher – A20008 |

**Table S4.**
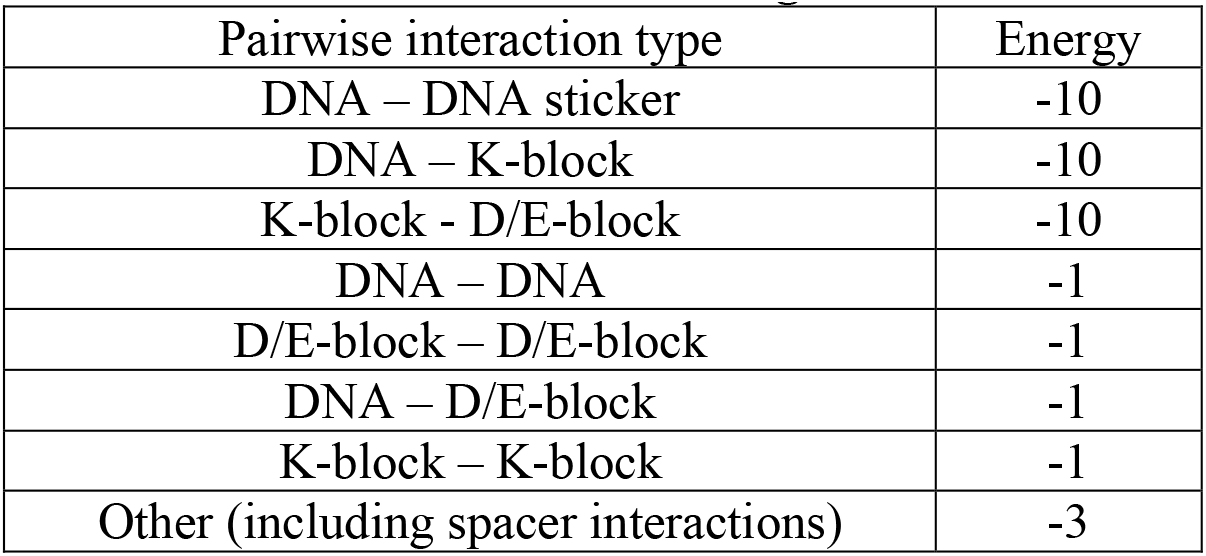
Pairwise interaction energies used in LaSSI simulations.

## References and Notes

(1) Shin, Y.; Brangwynne, C. P. Liquid Phase Condensation in Cell Physiology and Disease. Science 2017, 357 (6357), eaaf4382. https://doi.org/10.1126/science.aaf4382.

(2) Mittag, T.; Pappu, R. V. A Conceptual Framework for Understanding Phase Separation and Addressing Open Questions and Challenges. Mol. Cell 2022. https://doi.org/10.1016/j.molcel.2022.05.018.

(3) Banani, S. F.; Lee, H. O.; Hyman, A. A.; Rosen, M. K. Biomolecular Condensates: Organizers of Cellular Biochemistry. Nat. Rev. Mol. Cell Biol. 2017, 18 (5), 285–298. https://doi.org/10.1038/nrm.2017.7.

(4) Lafontaine, D. L. J.; Riback, J. A.; Bascetin, R.; Brangwynne, C. P. The Nucleolus as a Multiphase Liquid Condensate. Nat. Rev. Mol. Cell Biol. 2020, 1–18. https://doi.org/10.1038/s41580-020-0272-6.

(5) Riback, J. A.; Zhu, L.; Ferrolino, M. C.; Tolbert, M.; Mitrea, D. M.; Sanders, D. W.; Wei, M.-T.; Kriwacki, R. W.; Brangwynne, C. P. Composition-Dependent Thermodynamics of Intracellular Phase Separation. Nature 2020, 581 (7807), 209–214. https://doi.org/10.1038/s41586-020-2256-2.

(6) Pederson, T. The Nucleolus. Cold Spring Harb. Perspect. Biol. 2011, 3 (3), a000638. https://doi.org/10.1101/cshperspect.a000638.

(7) Feric, M.; Vaidya, N.; Harmon, T. S.; Mitrea, D. M.; Zhu, L.; Richardson, T. M.; Kriwacki, R. W.; Pappu, R. V.; Brangwynne, C. P. Coexisting Liquid Phases Underlie Nucleolar Subcompartments. Cell 2016, 165 (7), 1686–1697. https://doi.org/10.1016/j.cell.2016.04.047.

(8) Choi, J.-M.; Holehouse, A. S.; Pappu, R. V. Physical Principles Underlying the Complex Biology of Intracellular Phase Transitions. Annu. Rev. Biophys. 2020, 49 (1), 107–133. https://doi.org/10.1146/annurev-biophys-121219-081629.

(9) Wang, J.; Choi, J.-M.; Holehouse, A. S.; Lee, H. O.; Zhang, X.; Jahnel, M.; Maharana, S.; Lemaitre, R.; Pozniakovsky, A.; Drechsel, D.; Poser, I.; Pappu, R. V.; Alberti, S.; Hyman, A. A Molecular Grammar Governing the Driving Forces for Phase Separation of Prion-like RNA Binding Proteins. Cell 2018, 174 (3), 688-699.e16. https://doi.org/10.1016/j.cell.2018.06.006.

(10) Ruff, K. M.; Choi, Y. H.; Cox, D.; Ormsby, A. R.; Myung, Y.; Ascher, D. B.; Radford, S. E.; Pappu, R. V.; Hatters, D. M. Sequence Grammar Underlying the Unfolding and Phase Separation of Globular Proteins. Mol. Cell 2022. https://doi.org/10.1016/j.molcel.2022.06.024.

(11) Bremer, A.; Farag, M.; Borcherds, W. M.; Peran, I.; Martin, E. W.; Pappu, R. V.; Mittag, T. Deciphering How Naturally Occurring Sequence Features Impact the Phase Behaviours of Disordered Prion-like Domains. Nat. Chem. 2021, 1–12. https://doi.org/10.1038/s41557-021-00840-w.

(12) Greig, J. A.; Nguyen, T. A.; Lee, M.; Holehouse, A. S.; Posey, A. E.; Pappu, R. V.; Jedd, G. Arginine-Enriched Mixed-Charge Domains Provide Cohesion for Nuclear Speckle Condensation. Mol. Cell 2020. https://doi.org/10.1016/j.molcel.2020.01.025.

(13) Cohan, M. C.; Kyung Shinn, M.; Lalmansingh, J. M.; Pappu, R. V. Uncovering Non-Random Binary Patterns within Sequences of Intrinsically Disordered Proteins. J. Mol. Biol. 2021, 167373. https://doi.org/10.1016/j.jmb.2021.167373.

(14) Zarin, T.; Strome, B.; Peng, G.; Pritišanac, I.; Forman-Kay, J. D.; Moses, A. M. Identifying Molecular Features That Are Associated with Biological Function of Intrinsically Disordered Protein Regions. eLife 2021, 10, e60220. https://doi.org/10.7554/eLife.60220.

(15) Hubstenberger, A.; Courel, M.; Bénard, M.; Souquere, S.; Ernoult-Lange, M.; Chouaib, R.; Yi, Z.; Morlot, J.-B.; Munier, A.; Fradet, M.; Daunesse, M.; Bertrand, E.; Pierron, G.; Mozziconacci, J.; Kress, M.; Weil, D. P-Body Purification Reveals the Condensation of Repressed MRNA Regulons. Mol. Cell 2017, 68 (1), 144-157.e5. https://doi.org/10.1016/j.molcel.2017.09.003.

(16) Jain, S.; Wheeler, J. R.; Walters, R. W.; Agrawal, A.; Barsic, A.; Parker, R. ATPase-Modulated Stress Granules Contain a Diverse Proteome and Substructure. Cell 2016, 164 (3), 487–498. https://doi.org/10.1016/j.cell.2015.12.038.

(17) Thul, P. J.; Åkesson, L.; Wiking, M.; Mahdessian, D.; Geladaki, A.; Blal, H. A.; Alm, T.; Asplund, A.; Björk, L.; Breckels, L. M.; Bäckström, A.; Danielsson, F.; Fagerberg, L.; Fall, J.; Gatto, L.; Gnann, C.; Hober, S.; Hjelmare, M.; Johansson, F.; Lee, S.; Lindskog, C.; Mulder, J.; Mulvey, C. M.; Nilsson, P.; Oksvold, P.; Rockberg, J.; Schutten, R.; Schwenk, J. M.; Sivertsson, Å.; Sjöstedt, E.; Skogs, M.; Stadler, C.; Sullivan, D. P.; Tegel, H.; Winsnes, C.; Zhang, C.; Zwahlen, M.; Mardinoglu, A.; Pontén, F.; Feilitzen, K. von; Lilley, K. S.; Uhlén, M.; Lundberg, E. A Subcellular Map of the Human Proteome. Science 2017, 356 (6340). https://doi.org/10.1126/science.aal3321.

(18) Lee, B.; Jaberi-Lashkari, N.; Calo, E. A Unified View of Low Complexity Regions (LCRs) across Species. eLife 2022, 11, e77058. https://doi.org/10.7554/eLife.77058.

(19) Scott, M. S.; Boisvert, F.-M.; McDowall, M. D.; Lamond, A. I.; Barton, G. J. Characterization and Prediction of Protein Nucleolar Localization Sequences. Nucleic Acids Res. 2010, 38 (21), 7388–7399. https://doi.org/10.1093/nar/gkq653.

(20) Li, H.; Wang, B.; Yang, A.; Lu, R.; Wang, W.; Zhou, Y.; Shi, G.; Kwon, S. W.; Zhao, Y.; Jin, Y. Ly-1 Antibody Reactive Clone Is an Important Nucleolar Protein for Control of Self-Renewal and Differentiation in Embryonic Stem Cells. STEM CELLS 2009, 27 (6), 1244–1254. https://doi.org/10.1002/stem.55.

(21) Grob, A.; Colleran, C.; McStay, B. Construction of Synthetic Nucleoli in Human Cells Reveals How a Major Functional Nuclear Domain Is Formed and Propagated through Cell Division. Genes Dev. 2014, 28 (3), 220–230. https://doi.org/10.1101/gad.234591.113.

(22) Copenhaver, G. P.; Putnam, C. D.; Denton, M. L.; Pikaard, C. S. The RNA Polymerase I Transcription Factor UBF Is a Sequence-Tolerant HMG-Box Protein That Can Recognize Structured Nucleic Acids. Nucleic Acids Res. 1994, 22 (13), 2651–2657. https://doi.org/10.1093/nar/22.13.2651.

(23) Lai, Y.-S.; Tseng, H.-B.; Hu, C.-H. The Dimerization Domain of Upstream Binding Factor Contains Multiple Helical Structures. Biochem. Biophys. Res. Commun. 1996, 220 (3), 816– 823. https://doi.org/10.1006/bbrc.1996.0487.

(24) McStay, B.; Frazier, M. W.; Reeder, R. H. XUBF Contains a Novel Dimerization Domain Essential for RNA Polymerase I Transcription. Genes Dev. 1991, 5 (11), 1957–1968. https://doi.org/10.1101/gad.5.11.1957.

(25) Pak, C. W.; Kosno, M.; Holehouse, A. S.; Padrick, S. B.; Mittal, A.; Ali, R.; Yunus, A. A.; Liu, D. R.; Pappu, R. V.; Rosen, M. K. Sequence Determinants of Intracellular Phase Separation by Complex Coacervation of a Disordered Protein. Mol. Cell 2016, 63 (1), 72– 85. https://doi.org/10.1016/j.molcel.2016.05.042.

(26) Dinic, J.; Marciel, A. B.; Tirrell, M. V. Polyampholyte Physics: Liquid–Liquid Phase Separation and Biological Condensates. Curr. Opin. Colloid Interface Sci. 2021, 54, 101457. https://doi.org/10.1016/j.cocis.2021.101457.

(27) Stott, K.; Watson, M.; Bostock, M. J.; Mortensen, S. A.; Travers, A.; Grasser, K. D.; Thomas, J. O. Structural Insights into the Mechanism of Negative Regulation of Single-Box High Mobility Group Proteins by the Acidic Tail Domain *. J. Biol. Chem. 2014, 289 (43), 29817–29826. https://doi.org/10.1074/jbc.M114.591115.

(28) Choi, J.-M.; Dar, F.; Pappu, R. V. LASSI: A Lattice Model for Simulating Phase Transitions of Multivalent Proteins. PLOS Comput. Biol. 2019, 15 (10), e1007028. https://doi.org/10.1371/journal.pcbi.1007028.

(29) Tajrishi, M. M.; Tuteja, R.; Tuteja, N. Nucleolin. Commun. Integr. Biol. 2011, 4 (3), 267– 275. https://doi.org/10.4161/cib.4.3.14884.

(30) Okuwaki, M.; Saotome-Nakamura, A.; Yoshimura, M.; Saito, S.; Hirawake-Mogi, H.; Sekiya, T.; Nagata, K. RNA-Recognition Motifs and Glycine and Arginine-Rich Region Cooperatively Regulate the Nucleolar Localization of Nucleolin. J. Biochem. (Tokyo) 2021, 169 (1), 87–100. https://doi.org/10.1093/jb/mvaa095.

(31) Mitrea, D. M.; Cika, J. A.; Stanley, C. B.; Nourse, A.; Onuchic, P. L.; Banerjee, P. R.; Phillips, A. H.; Park, C.-G.; Deniz, A. A.; Kriwacki, R. W. Self-Interaction of NPM1 Modulates Multiple Mechanisms of Liquid–Liquid Phase Separation. Nat. Commun. 2018, 9 (1), 842. https://doi.org/10.1038/s41467-018-03255-3.

(32) Banerjee, P. R.; Milin, A. N.; Moosa, M. M.; Onuchic, P. L.; Deniz, A. A. Reentrant Phase Transition Drives Dynamic Substructure Formation in Ribonucleoprotein Droplets. Angew. Chem. Int. Ed. 2017, 56 (38), 11354–11359. https://doi.org/10.1002/anie.201703191.

(33) Henninger, J. E.; Oksuz, O.; Shrinivas, K.; Sagi, I.; LeRoy, G.; Zheng, M. M.; Andrews, J. O.; Zamudio, A. V.; Lazaris, C.; Hannett, N. M.; Lee, T. I.; Sharp, P. A.; Cissé, I. I.; Chakraborty, A. K.; Young, R. A. RNA-Mediated Feedback Control of Transcriptional Condensates. Cell 2020. https://doi.org/10.1016/j.cell.2020.11.030.

(34) Wu, M.; Xu, G.; Han, C.; Luan, P.-F.; Xing, Y.-H.; Nan, F.; Yang, L.-Z.; Huang, Y.; Yang, Z.-H.; Shan, L.; Yang, L.; Liu, J.; Chen, L.-L. LncRNA SLERT Controls Phase Separation of FC/DFCs to Facilitate Pol I Transcription. Science 2021, 373 (6554), 547–555. https://doi.org/10.1126/science.abf6582.

## References

(1) The UniProt Consortium. UniProt: The Universal Protein Knowledgebase in 2021. Nucleic Acids Res. 2021, 49 (D1), D480–D489. https://doi.org/10.1093/nar/gkaa1100.

(2) Piovesan, D.; Necci, M.; Escobedo, N.; Monzon, A. M.; Hatos, A.; Micetic, I.; Quaglia, F.; Paladin, L.; Ramasamy, P.; Dosztányi, Z.; Vranken, W. F.; Davey, N. E.; Parisi, G.; Fuxreiter, M.; Tosatto, S. C. E. MobiDB: Intrinsically Disordered Proteins in 2021. Nucleic Acids Res. 2021, 49 (D1), D361–D367. https://doi.org/10.1093/nar/gkaa1058.

(3) Zarin, T.; Strome, B.; Nguyen Ba, A. N.; Alberti, S.; Forman-Kay, J. D.; Moses, A. M. Proteome-Wide Signatures of Function in Highly Diverged Intrinsically Disordered Regions. eLife 2019, 8, e46883. https://doi.org/10.7554/eLife.46883.

(4) Cohan, M. C.; Kyung Shinn, M.; Lalmansingh, J. M.; Pappu, R. V. Uncovering Non-Random Binary Patterns within Sequences of Intrinsically Disordered Proteins. J. Mol. Biol. 2021, 167373. https://doi.org/10.1016/j.jmb.2021.167373.

(5) Holehouse, A. S.; Das, R. K.; Ahad, J. N.; Richardson, M. O. G.; Pappu, R. V. CIDER: Resources to Analyze Sequence-Ensemble Relationships of Intrinsically Disordered Proteins. Biophys. J. 2017, 112 (1), 16–21. https://doi.org/10.1016/j.bpj.2016.11.3200.

(6) Thul, P. J.; Åkesson, L.; Wiking, M.; Mahdessian, D.; Geladaki, A.; Blal, H. A.; Alm, T.; Asplund, A.; Björk, L.; Breckels, L. M.; Bäckström, A.; Danielsson, F.; Fagerberg, L.; Fall, J.; Gatto, L.; Gnann, C.; Hober, S.; Hjelmare, M.; Johansson, F.; Lee, S.; Lindskog, C.; Mulder, J.; Mulvey, C. M.; Nilsson, P.; Oksvold, P.; Rockberg, J.; Schutten, R.; Schwenk, J. M.; Sivertsson, Å.; Sjöstedt, E.; Skogs, M.; Stadler, C.; Sullivan, D. P.; Tegel, H.; Winsnes, C.; Zhang, C.; Zwahlen, M.; Mardinoglu, A.; Pontén, F.; Feilitzen, K. von; Lilley, K. S.; Uhlén, M.; Lundberg, E. A Subcellular Map of the Human Proteome. Science 2017, 356 (6340). https://doi.org/10.1126/science.aal3321.

(7) Jain, S.; Wheeler, J. R.; Walters, R. W.; Agrawal, A.; Barsic, A.; Parker, R. ATPase-Modulated Stress Granules Contain a Diverse Proteome and Substructure. Cell 2016, 164 (3), 487–498. https://doi.org/10.1016/j.cell.2015.12.038.

(8) Hubstenberger, A.; Courel, M.; Bénard, M.; Souquere, S.; Ernoult-Lange, M.; Chouaib, R.; Yi, Z.; Morlot, J.-B.; Munier, A.; Fradet, M.; Daunesse, M.; Bertrand, E.; Pierron, G.; Mozziconacci, J.; Kress, M.; Weil, D. P-Body Purification Reveals the Condensation of Repressed MRNA Regulons. Mol. Cell 2017, 68 (1), 144-157.e5. https://doi.org/10.1016/j.molcel.2017.09.003.

(9) Nagaraj, N.; Wisniewski, J. R.; Geiger, T.; Cox, J.; Kircher, M.; Kelso, J.; Pääbo, S.; Mann, M. Deep Proteome and Transcriptome Mapping of a Human Cancer Cell Line. Mol. Syst. Biol. 2011, 7 (1), 548. https://doi.org/10.1038/msb.2011.81.

(10) Ouyang, W.; Winsnes, C. F.; Hjelmare, M.; Cesnik, A. J.; Åkesson, L.; Xu, H.; Sullivan, D. P.; Dai, S.; Lan, J.; Jinmo, P.; Galib, S. M.; Henkel, C.; Hwang, K.; Poplavskiy, D.; Tunguz, B.; Wolfinger, R. D.; Gu, Y.; Li, C.; Xie, J.; Buslov, D.; Fironov, S.; Kiselev, A.; Panchenko, D.; Cao, X.; Wei, R.; Wu, Y.; Zhu, X.; Tseng, K.-L.; Gao, Z.; Ju, C.; Yi, X.; Zheng, H.; Kappel, C.; Lundberg, E. Analysis of the Human Protein Atlas Image Classification Competition. Nat. Methods 2019, 16 (12), 1254–1261. https://doi.org/10.1038/s41592-019-0658-6.

(11) Ouyang, W.; Mueller, F.; Hjelmare, M.; Lundberg, E.; Zimmer, C. ImJoy: An Open-Source Computational Platform for the Deep Learning Era. Nat. Methods 2019, 16 (12), 1199– 1200. https://doi.org/10.1038/s41592-019-0627-0.

(12) Choi, J.-M.; Dar, F.; Pappu, R. V. LASSI: A Lattice Model for Simulating Phase Transitions of Multivalent Proteins. PLOS Comput. Biol. 2019, 15 (10), e1007028. https://doi.org/10.1371/journal.pcbi.1007028.

(13) Farag, M.; Cohen, S. R.; Borcherds, W. M.; Bremer, A.; Mittag, T.; Pappu, R. V. Condensates of Disordered Proteins Have Small-World Network Structures and Interfaces Defined by Expanded Conformations. bioRxiv May 26, 2022, p 2022.05.21.492916. https://doi.org/10.1101/2022.05.21.492916.

(14) Feric, M.; Vaidya, N.; Harmon, T. S.; Mitrea, D. M.; Zhu, L.; Richardson, T. M.; Kriwacki, R. W.; Pappu, R. V.; Brangwynne, C. P. Coexisting Liquid Phases Underlie Nucleolar Subcompartments. Cell 2016, 165 (7), 1686–1697. https://doi.org/10.1016/j.cell.2016.04.047.

(15) Zhang, H.; Elbaum-Garfinkle, S.; Langdon, E. M.; Taylor, N.; Occhipinti, P.; Bridges, A. A.; Brangwynne, C. P.; Gladfelter, A. S. RNA Controls PolyQ Protein Phase Transitions. Mol. Cell 2015, 60 (2), 220–230. https://doi.org/10.1016/j.molcel.2015.09.017.

